# Polycystins recruit cargo to distinct ciliary extracellular vesicle subtypes

**DOI:** 10.1101/2024.04.17.588758

**Authors:** Inna A. Nikonorova, Elizabeth desRanleau, Katherine C. Jacobs, Joshua Saul, Jonathon D. Walsh, Juan Wang, Maureen M. Barr

## Abstract

Therapeutic use of tiny extracellular vesicles (EVs) requires understanding cargo loading mechanisms. Here, we used a modular proximity label approach to identify EV cargo associated with the transient potential channel (TRP) polycystin PKD-2 of *C. elegans*. Polycystins are conserved receptor-TRP channel proteins affecting cilium function; dysfunction causes polycystic kidney disease in humans and mating deficits in *C. elegans*. Polycystin-2 EV localization is conserved from algae to humans, hinting at an ancient and unknown function. We discovered that polycystins associate with and direct specific cargo to EVs: channel-like PACL-1, dorsal and ventral membrane C-type lectins PAMLs, and conserved tumor necrosis-associated factor (TRAF) signaling adaptors TRF-1 and TRF-2. Loading of these components relied on polycystin-1 LOV-1. Our modular EV-TurboID approach can be applied in both cell– and tissue-specific manners to define the composition of distinct EV subtypes, addressing a major challenge of the EV field.

## Introduction

Studying extracellular vesicles (EVs) presents significant challenges due to their diverse types, sizes, and compositions. This heterogeneity complicates isolation, characterization, and analysis. Addressing the challenge of cargo composition of individual EV subtypes in the face of EV diversity, however, is crucial for advancing EV-based diagnostic, therapeutic, and biomedical research.

Here we introduce a methodology to identify the cargo of individual EV subtypes via proximity labeling in living *C. elegans*. We took this approach to pinpoint the cargo carried by specific ciliary EV subtypes carrying the transient receptor potential (TRP) polycystin channel PKD-2. Human PKD2 and PKD1 are mutated in autosomal dominant polycystic kidney disease, the most common inherited cause of renal failure^1^. PKD2 and PKD1 encode a TRP polycystin channel and receptor complex that function in the primary cilia of kidney tubules and are cargo of urinary EVs^2^.

Polycystins are highly conserved and function in vertebrate development and homeostasis, with roles ranging from kidney tubule function, spinal cord morphogenesis to establishing left–right axis patterning. In the morphogenesis of the spinal cord, PKD2L1 functions in CSF-contacting neurons that are critical for precise posture control^3–5^. During gastrulation, polycystin-2 and PKD1L1 (polycystic kidney disease 1 – like 1) establish laterality, possibly via EV-like particles ^6^.

In *C. elegans*, the polycystins LOV-1 and PKD-2 co-localize in cilia of male-specific sensory neurons and are required for multiple aspects of male sexual behaviors^7–11^. LOV-1 and PKD-2 also colocalize on ciliary EVs released from male-specific ciliated neurons and are shed from the cilium’s tip into the environment ^12^. These polycystin-carrying EVs are transferred from the male to the hermaphrodite during mating and may facilitate inter-animal communication ^13,14^. Polycystin EV localization is conserved from algae^15^ to *C. elegans* to humans, hinting at an ancient but, as of yet, unknown function.

Akin to other systems, *C. elegans* EVs share similar size and density, despite diverse biogenesis routes and cell sources. We recently isolated *C. elegans* EVs in density gradients guided by the presence of fluorescence-labeled polycystin-2 PKD-2::GFP and showed that many other EV subtypes co-isolate with polycystin-carrying EVs ^16^. Herein, we developed a system for precise identification of cargo of individual EV subtypes and applied this EV-TurboID methodology to identify and validate PKD-2 interactors in cilia and ciliary EVs of *C. elegans*.

## Results

### Identification of polycystin proximity interactors within EVs

A critical objective in deciphering in vivo EV functions is defining EV subtypes and identifying their cargo. Thus, we sought to exploit features of the *C. elegans* model for protein identification. We engineered proximity labeling enzyme TurboID^17^ to be targeted to a GFP-labeled ciliary EV cargo PKD-2::GFP via a GFP-binding nanobody domain^18^. Between the GFP-binding nanobody and TurboID domains, we added a fluorescent mScarlet domain that would enable us to track TurboID in living animals and environmentally released EVs through super-resolution live imaging (Fig. 1a-c). We observed anti-GFP-nanobody::mScarlet::TurboID and PKD-2::GFP colocalization in male-specific neuronal cell bodies, cilia, and environmentally released ciliary EVs (Fig. 1c, Extended Data Fig. 1a-b).

**Fig. 1.**
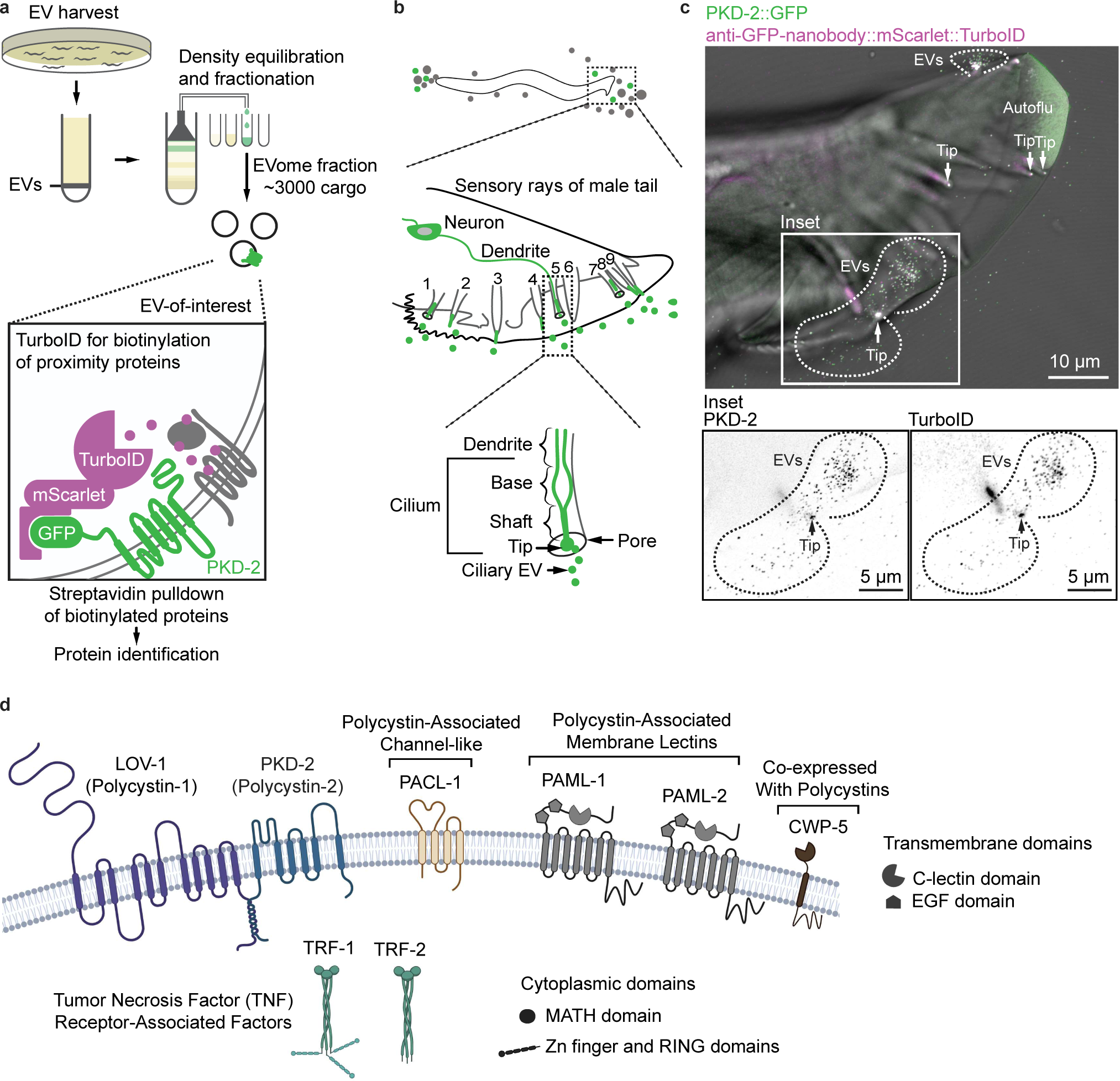
Proximity labeling within PKD-2 EVs identified candidate interactors. **a,** Scheme of EV harvest and enrichment, followed by pulldown of candidate interactors biotinylated by TurboID targeted to PKD-2 EVs. **b,** Scheme of male tail anatomical structures that release PKD-2 EVs. Each sensory ray of the male tail harbors a ciliated dendritic ending protruding into the environment and releasing PKD-2 EVs from the tip of its sensory cilium. **c,** Visualization of PKD-2::GFP EVs carrying TurboID. **d,** Identified top candidate interactors.

To identify EV cargo proteins in the proximity of the targeted polycystin, we pulled down biotinylated proteins for mass spectrometry analysis from a density gradient fraction enriched in PKD-2::GFP EVs (Fig. 1a, d, Table S1). Proximity interactors included two known cargo of PKD-2::GFP: LOV-1^7–9^ and the ciliary EV biogenesis protein CIL-7^14,19^, confirming the efficacy of our EV-TurboID approach. We also identified and validated six novel cargo: two homologs of human TNF receptor-associated factors (TRAFs) TRF-1 and TRF-2, three transmembrane lectins (PAML-1, PAML-2, CWP-5), and a novel transmembrane protein resembling an ion channel (PACL-1) (Fig. 1d; for the complete list of identified proteins, refer to Table S1). *trf-1*, *trf-2*, and *cwp-5* mutants are defective in polycystin-mediated mating behaviors^20–22^. *trf-1*, *trf-2*, *paml-2*, and *cwp-5* are co-expressed with *pkd-2*^20,21^. further demonstration of the efficacy of EV-TurboID. The latter experiments were performed through overexpression of transgenes from extrachromosomal arrays, and their protein localization in the cilium and/or EVs had not been determined due to limitations in epifluorescence microscopy used at the time. In this study, we took an unbiased validation approach by generating endogenous reporters for all top candidates and PKD-2 using CRISPR/Cas9-mediated genome editing. We used super-resolution microscopy to analyze candidate co-localization with PKD-2 within cilia and ciliary EVs (Fig. 2a).

**Fig. 2.**
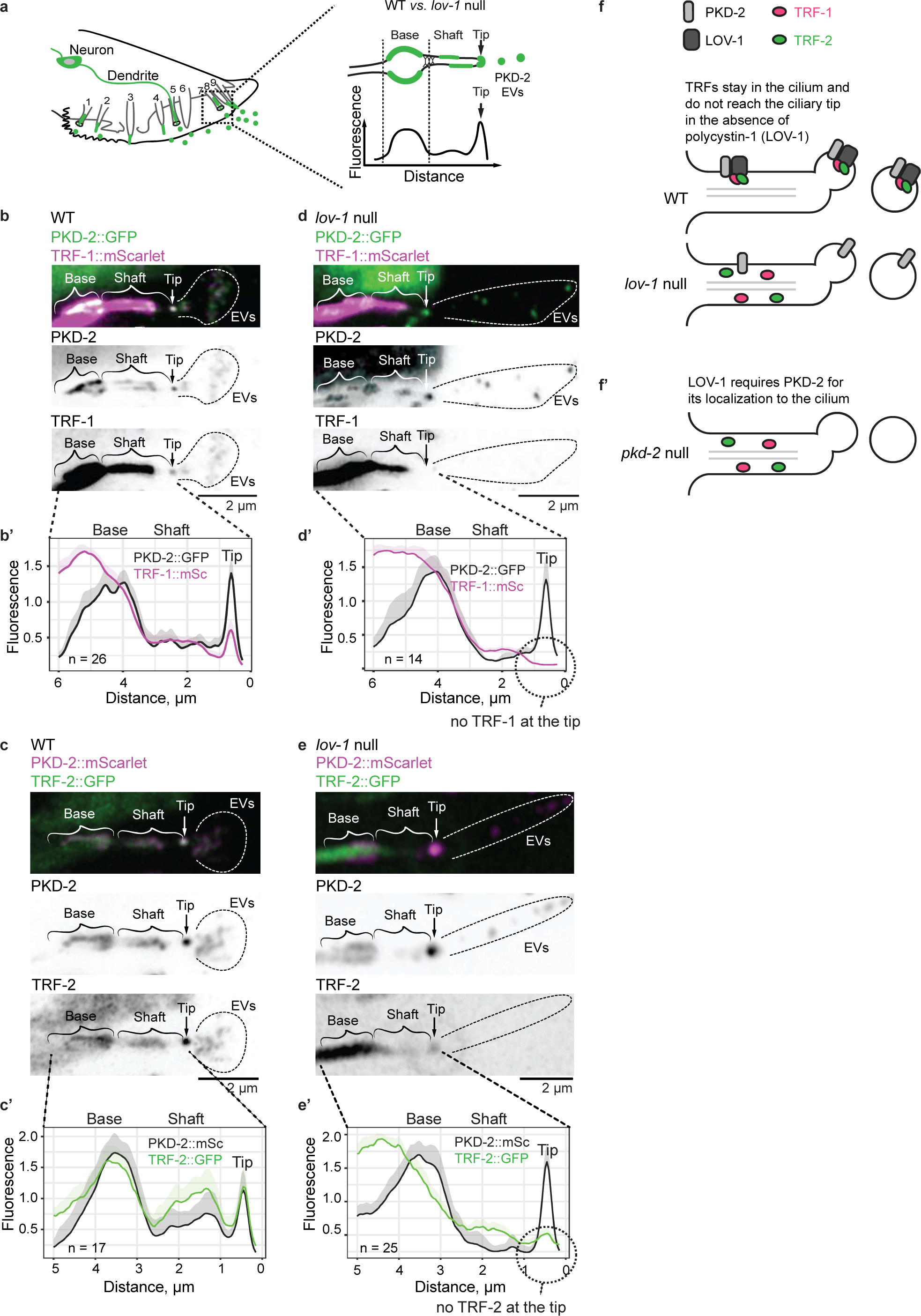
LOV-1 is required for loading TRFs to ciliary PKD-2 EVs. **a,** Scheme of fluorescent profiling along cilia. **b-e,** Flattened z-stacks show the ciliary presence of endogenous FP-tagged PKD-2 with TRF-1::mScarlet (**b**) and TRF-2::GFP (**c**) in the wild-type cilium and *lov-1* mutant (**d, e**). Representative average fluorescent profiles for each case are shown (**b’, c’, d’, e’**). Fluorescence values are normalized to the average of minimum and maximum values for each cilium. **f-f’,** Scheme of the molecular mechanism for loading TRFs to ciliary EVs. In the *lov-1* mutant, cilia produce EVs without TRFs (**f**), whereas in the *pkd-2* mutant, ciliary EVs contain neither polycystins nor TRFs (**f’**).

### Polycystin EVs carry homologs of human TRAFs – TRFs

The TNF receptor-associated factor (TRAFs) family comprises adapter-type proteins that act downstream of receptors and mediate the assembly of cytoplasmic signaling components^23^. We discovered that endogenous *trf-1* and *trf-2* genes are expressed at high levels in polycystin-expressing sensory neurons of males important for male sexual behaviors and mating processes (<u>c</u>ephalic <u>m</u>ale CEMs, tail <u>R</u>ay <u>B</u> neurons 1-5, 7-9, and <u>ho</u>ok sensillum B <u>n</u>euron HOB). In these neurons, TRF-1 and TRF-2 colocalized with PKD-2 in cilia and EVs (Fig. 2b – c’, Extended Data Fig. 2a-c), consistent with TRAFs acting as adaptors for the polycystin complex and with polycystin EVs serving as carriers for TRAFs to engage target tissues.

### Polycystins load TRFs to ciliary EVs

To test whether polycystins associate with TRAFs *in vivo*, we analyzed TRAF subcellular localization in the background of disrupted polycystins. LOV-1 has an extensive N-terminal domain that might act as a receptor for the polycystin complex. We previously showed that PKD-2 is essential to bring LOV-1 to the cilium and EVs, while *lov-1* is not required for PKD-2 ciliary and EV localization^9^. Therefore, we examined the localization of the TRAFs and other EV cargo candidates in the absence of *lov-1*. In the *lov-1* mutant, TRF-1 and TRF-2 remained in the ciliary shaft, failed to reach the ciliary tip, and were not loaded and released in ciliary EVs (Fig. 2d-f, Extended Data Fig. 2a-c). These data indicate that LOV-1 is required for loading TRF-1 and TRF-2 to ciliary EVs, but is not required for their localization to the cilium shaft.

Mutations in polycystins lead to a failure in *C. elegans* male mating^14,19^. Similar behavioral deficiencies are exhibited by *trf-1* and *trf-2* mutants, indicating that TRAF and polycystins act in the same genetic pathway ^20^. Despite this, disrupting *trf-1* and *trf-2* genes did not affect PKD-2 and LOV-1 co-localization in cilia or their release in EVs (Extended Data Fig. 4a-b), indicating that TRF-1 and TRF-2 are non-essential for PKD-2 and LOV-1 ciliary and EV localization. Taken together these data suggest that TRF-1 and TRF-2 may act as signaling adaptors for the polycystins in the cilium to regulate male mating behavior.

### TRFs require each other for their loading to ciliary EVs

We next investigated the relationship between TRF-1 and TRF-2 in cilia and EVs. In the *trf-2* mutant, TRF-1::mScarlet remained confined to the ciliary shaft and failed to locate to ciliary EVs (Fig. 3a-a’, Extended Data Fig. 4a-b). Conversely, disruption of *trf-1* resulted in the absence of TRF-2::GFP in the ciliary shaft altogether (Fig. 3b-b’), indicating that TRF-1 is required for the entry of TRF-2 into the cilium. Additionally, TRF-2::GFP abundance was reduced in dendrites and accompanied by a three-fold reduction in cytoplasm, suggesting that TRF-1 might stabilize TRF-2 or shield it from degradation (Fig. 3c-c’, Extended Data Fig. 4c).

**Fig. 3.**
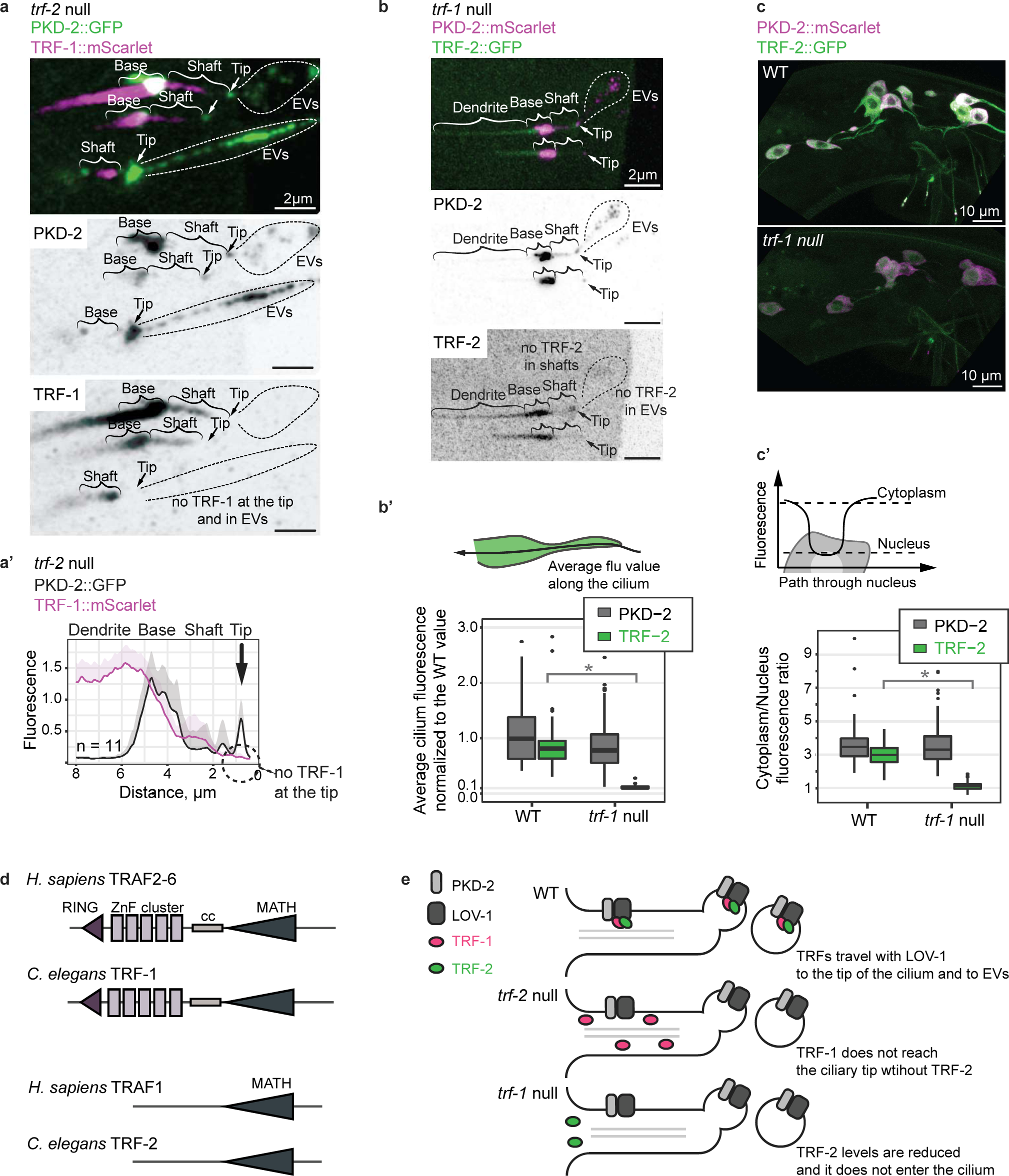
TRFs require each other for their loading to ciliary EVs. **a,** Flattened z-stack shows that disruption of *trf-2* abrogates loading of TRF-1::mScarlet to PKD-2::GFP EVs. TRF-1::mScarlet stays in the cilium and does not reach the ciliary tip, as shown by fluorescent profiling (**a’**). Fluorescence values are normalized to the average of minimum and maximum values for each cilium. **b,** Flattened z-stack shows that disruption of *trf-1* abrogates ciliary localization of TRF-2::GFP, and thus, no TRF-2::GFP is loaded to PKD-2::mScarlet EVs. **b’,** Quantification of total fluorescence within the cilium shows that TRF-2::GFP ciliary levels in the *trf-1* mutant are reduced tenfold compared to WT cilia, whereas PKD-2::mScarlet total levels are not affected (WT n = 17 animals and 73 cilia, *trf-1* null n = 36 animals and 112 cilia).. **c,** Flattened z-stack through cell bodies of RnB neurons showing reduced cytoplasmic levels of TRF-2::GFP, with quantification shown on panel **c’** (WT n = 11 animals and 96 cilia, *trf-1* null n = 15 animals and 147 cilia). **d,** Homology of *C. elegans* TRFs to human TRAFs. **e,** Scheme of the molecular mechanism for loading TRFs to ciliary EVs. In the *trf-2* mutant, cilia produce EVs without TRF-1, whereas the *trf-1* mutation causes reduced levels of TRF-2 and abrogates TRF-2 ciliary localization. *Mann-Whitney U Test *p* < .00001.

Evolutionary conservation of TRAF domain assemblies suggests that the function of human and *C. elegans* TRAFs might likewise be conserved. Both human and *C. elegans* TRAF families comprise two types of domain organizations: RING domain-containing TRAFs (TRF-1 in *C. elegans* and TRAF2-6 in humans) and RING-less TRAFs (TRF-2 in *C. elegans* and TRAF1 in humans) (Fig. 3d). The RING domain confers ubiquitin ligase activity to human TRAF6^24,25^, promoting the assembly of multimeric signaling networks presumably comprised of trimer dimers^26^. In *Dictyostelium discoideum,* the RING domain recruits TRAF to sites of membrane damage^27^. However, the specific role of RING-less TRAFs in the context of TRAF cascades remains unclear. This study shows that TRAFs act in the cilium (Fig. 3e). Specifically, our work highlights the role of TRF-1 (a RING-containing TRAF, likely functioning as a ubiquitin ligase) in ensuring the stability of TRF-2 and facilitating its entry into the cilium. Conversely, TRF-2 is essential for recruiting TRF-1 to ciliary EVs.

### Polycystin complex associates with and recruits a channel-like protein PACL-1 to cilia and EVs

Our proximity labeling studies also identified a novel four-pass transmembrane protein, T13F3.7, which we named PACL-1 (<u>P</u>olycystin-<u>A</u>ssociated <u>C</u>hannel-like protein) (Extended Data Fig. 5). To investigate whether PACL-1 functions with polycystins, we generated an endogenous fluorescently tagged reporter of PACL-1 in which GFP was linked to the intracellular C-terminus. The *pacl-1* gene was expressed specifically in all polycystin-expressing neurons. PACL-1::GFP colocalized with PKD-2::mScarlet in cilia and EVs (Fig. 4a, Extended Data Fig. 6a-b). This subcellular localization of PACL-1::GFP was disrupted by the *lov-1* mutation (Fig. 4b). In *lov-1* mutant males, PACL-1::GFP remained in neuronal cell bodies and was not localized to cilia or environmentally released EVs, indicating that LOV-1 recruits the PACL-1 channel-like protein to cilia and EVs (Fig. 4c).

**Fig. 4.**
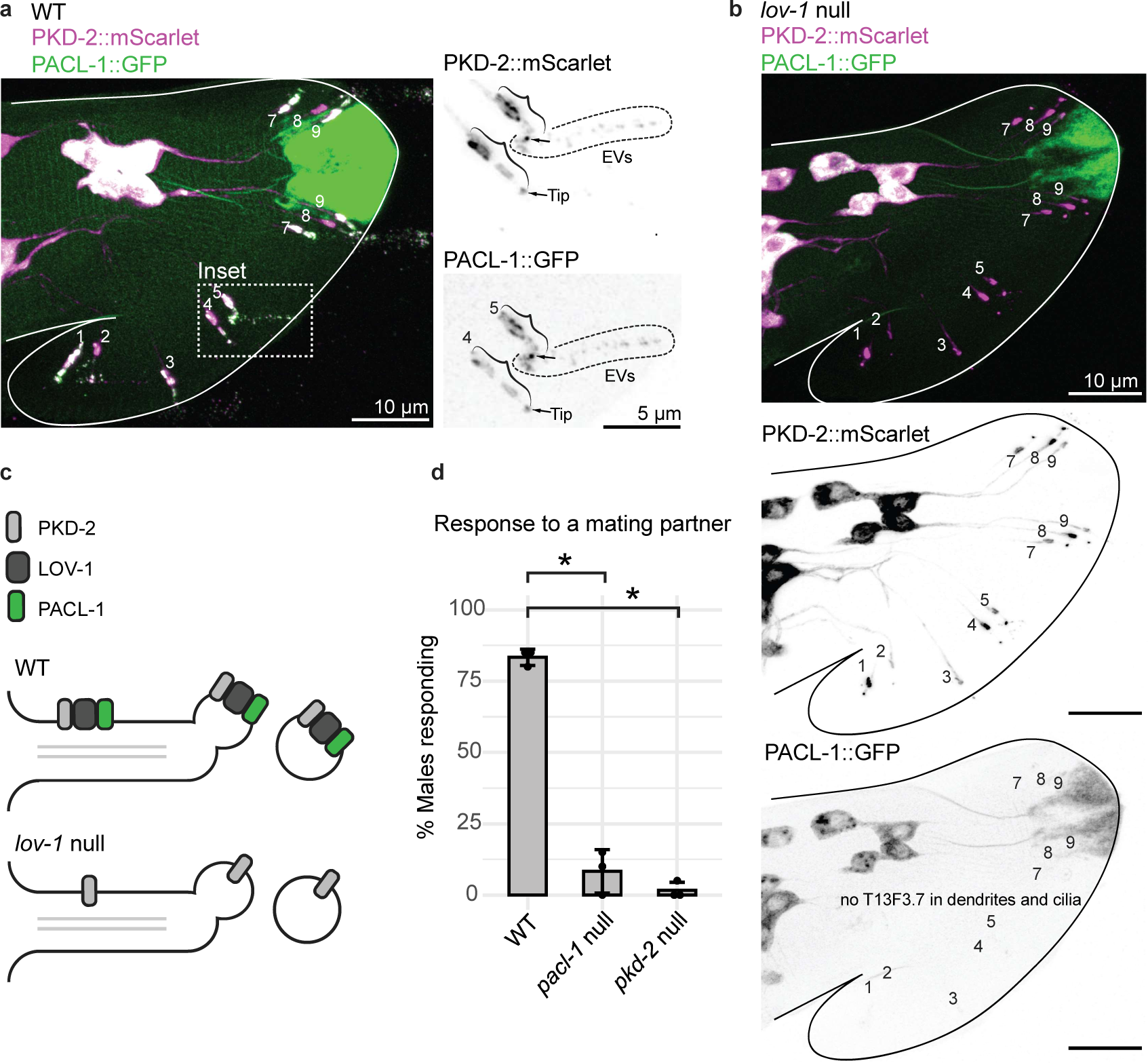
The polycystin complex associates with and recruits a channel-like protein PACL-1 and to cilia and EVs. **a-b,** Flattened z-stacks showing colocalization of PKD-2::mScarlet with PACL-1::GFP in WT (**a**) and *lov-1* mutant (**b**) animals. **c,** Scheme of molecular mechanism showing that loading of PACL-1::GFP to ciliary EVs requires LOV-1. **d,** Assessment of male mating behavior shows that the *pacl-1* mutant is deficient in responding to hermaphrodite contact. *Student’s t test p<0.05.

We then tested whether PACL-1 functions within the polycystin pathway by assessing behaviors regulated by polycystins in male-specific ciliated neurons – initiation of mating (response to hermaphrodite contact). The *pacl-1* mutant males were deficient in the response behavior (Fig. 4d), consistent with PACL-1 and the polycystins acting in the same sensory pathway. These data also suggest that the polycystin pathway uses an auxiliary ion channel associated with cilia and EVs to influence behavior.

### The polycystin complex associates with distinct transmembrane C-type lectins that specify dorsal and ventral populations of cilia and EVs

Our proximity labeling studies also identified novel multi-pass transmembrane Y70G10A.2 and F25D7.5 candidates, which possessed N-terminal extracellular EGF and C-type lectin domains (Fig. 1d). On the basis of these domains, we named these candidates <u>P</u>olycystin-<u>A</u>ssociated <u>M</u>embrane <u>L</u>ectins PAML-1 and PAML-2, respectively.

We generated endogenous reporters of PAML-1 and PAML-2. The *paml-1::gfp* expression was limited to a ventral subset of *pkd-2*-expressing neurons, specifically those in which cilia protrude to the ventral side of the mail tail – HOB, R2B, R3B, R4B, and R8B neurons. These ventrally located neurons mediate vulva location behavior and ventral contact-based response^28^. Within these ventral tail neurons, PAML-1::GFP colocalized with PKD-2::mScarlet in both cilia and EVs (Fig. 5a, Extended Data 7a-b).

**Fig. 5.**
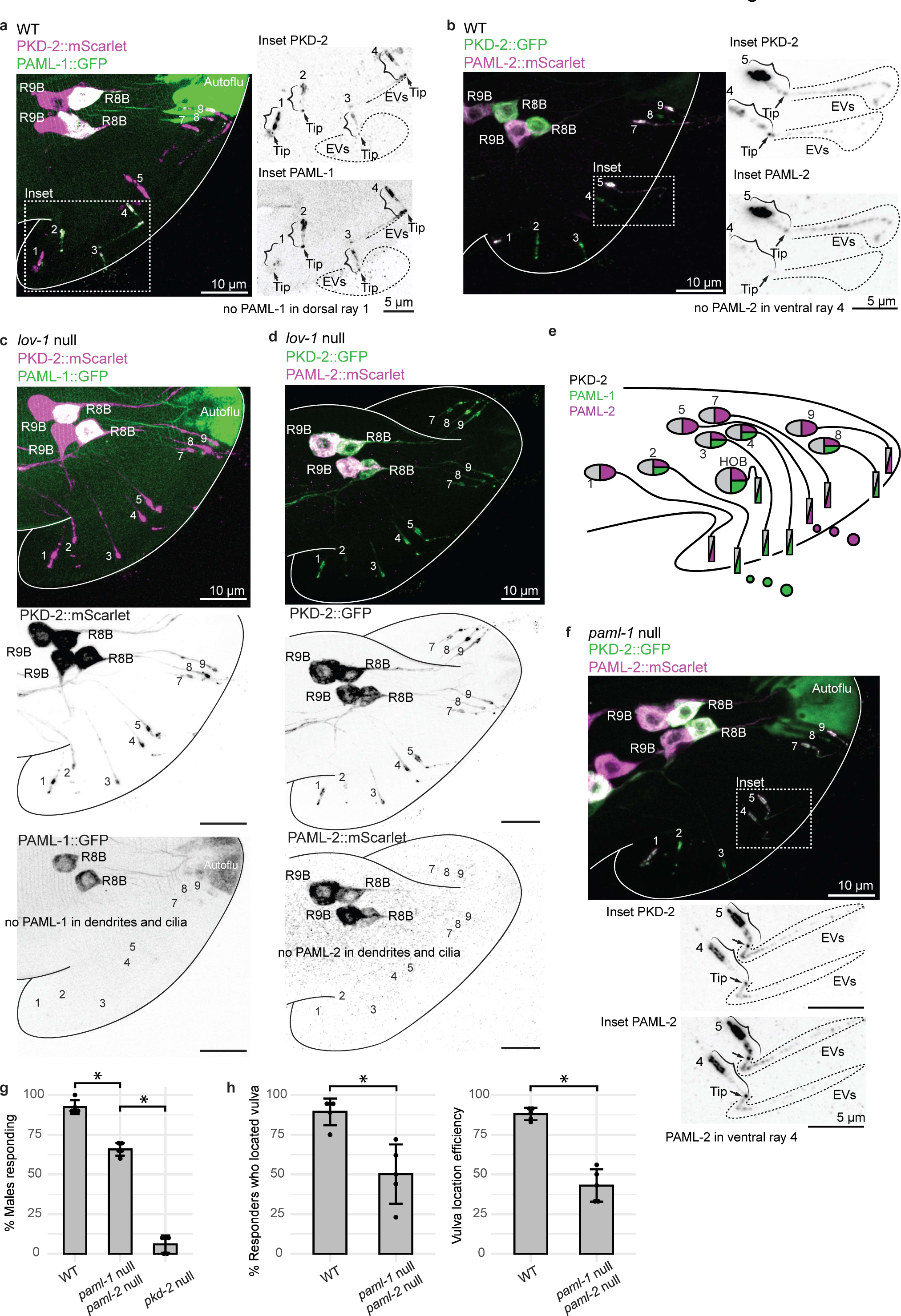
The polycystin complex associates with transmembrane C-type lectins that specify dorsal and ventral populations of polycystin cilia and EVs. **a-b,** Flattened z-stacks showing colocalization of PKD-2 reporters with PAML-1::GFP (**a**) and PAML-2::mScarlet (**b**) in WT. **c-d**, Disruption of *lov-1* abrogates ciliary localization of PAML-1::GFP (**c**) and PAML-2::mScarlet (**d**). **e,** Scheme of PAML-1::GFP and PAML-2::mScarlet localization in RnB neuronal cell bodies and cilia. **f,** Flattened z-stack showing that disruption of *pacl-1* results in PAML-2::mScarlet ciliary localization in ventral neurons. **g-h,** Assessment of male mating behavior shows that the *paml-1; paml-2* double mutant is deficient in responding to hermaphrodite contact (**g**), and location of vulva (**h**).* *Student’s t test p<0.05.

In contrast to *paml-1*, *paml-2* was expressed in all polycystin-expressing neurons, including HOB, Ray B neurons 1-5, 7-9 in the tail, and CEM neurons in the head (Extended Data Fig. 7c-d). However, PAML-2::mScarlet localized to cilia and EVs solely in the dorsal rays of the male tail (R1B, R5B, R7B, and R9B) (Fig. 5a-b, Extended Data 7c-d) and was absent from the PAML-1-carrying ventral ray cilia and EVs. Dorsally located ray neurons mediate dorsal contact-based response^28^.

The presence of PAML-1 and PAML-2 in cilia depended on LOV-1. In *lov-1* mutants, both PAMLs remained confined to the cell body and did not localize to dendrites, cilia, or ciliary EVs (Fig. 5c-d, Extended Data Fig. 8a-b). This suggests that LOV-1 recruits PAMLs to ciliary EVs and that this recruitment takes place in the cell body of the neuron. Here LOV-1 acts as a pivotal scaffold protein recruiting transmembrane components of dorso-ventral specialization to the cilium and polycystin-carrying EVs.

To uncover mechanisms behind the selective presence of PAML-2 solely in dorsal cilia (Fig. 5e), we examined the localization of PAML-2::mScarlet in *paml-1* mutant animals. In the absence of ventral *paml-1* expression, PAML-2::mScarlet ectopically localized to the cilia of ventral ray neurons (Fig. 5f). The PAML-2 total abundance in ventral cilia of *paml-1* mutants constituted 25-60% of the levels observed in dorsal neurons (Extended Data Fig. 8c). The ciliary presence of dorsal lectin PAML-2::mScarlet in ventral neurons of *paml-1* mutants was accompanied by a 2-4-fold increase in PAML-2::mScarlet abundance in the cell bodies of ventral neurons (Extended Data Fig. 8c’), suggesting compensatory expression mechanism that boosts production of PAML-2 in ventral neurons in the absence of the PAML-1 ventral lectin.

We then tested whether PAMLs are involved in the polycystin-mediated male mating behavior. The *paml-1; paml-2* double null mutant exhibited reduced response to hermaphrodite contact compared to wild-type animals (Fig. 5g). The *paml-1; paml-2* mutant response defect was not as severe as of the *pkd-2* mutant. The *paml-1; paml-2* double mutants also displayed defects in vulva location behavior (Fig. 5h). Taken together, our data suggest that PAML-1 and PAML-2 are recruited by polycystins to cilia and EVs in a specific dorsal-ventral pattern and act in sensory pathways (response and vulva location behaviors) mediated by polycystins.

### Polycystin complex is not required for shedding of ciliary EVs as evidenced by tracking cargo CWP-5

The final validated candidate interactor of PKD-2 was CWP-5 (<u>C</u>oexpressed <u>W</u>ith <u>P</u>olycystins 5), a single-pass transmembrane protein^21^ that functions in the polycystin-mediated male mating response to the hermaphrodite^22^. We generated the endogenous reporter CWP-5::GFP and found *cwp-5::gfp* expression in all *pkd-2*-expressing neuronal cell bodies except R2B (Extended Data Fig. 9a-b). The abundant ciliary localization of CWP-5::GFP was observed in all but R2B and R8B neurons (Fig. 6a, Extended Data Fig. 9c-c’, 10a-b).

**Fig. 6.**
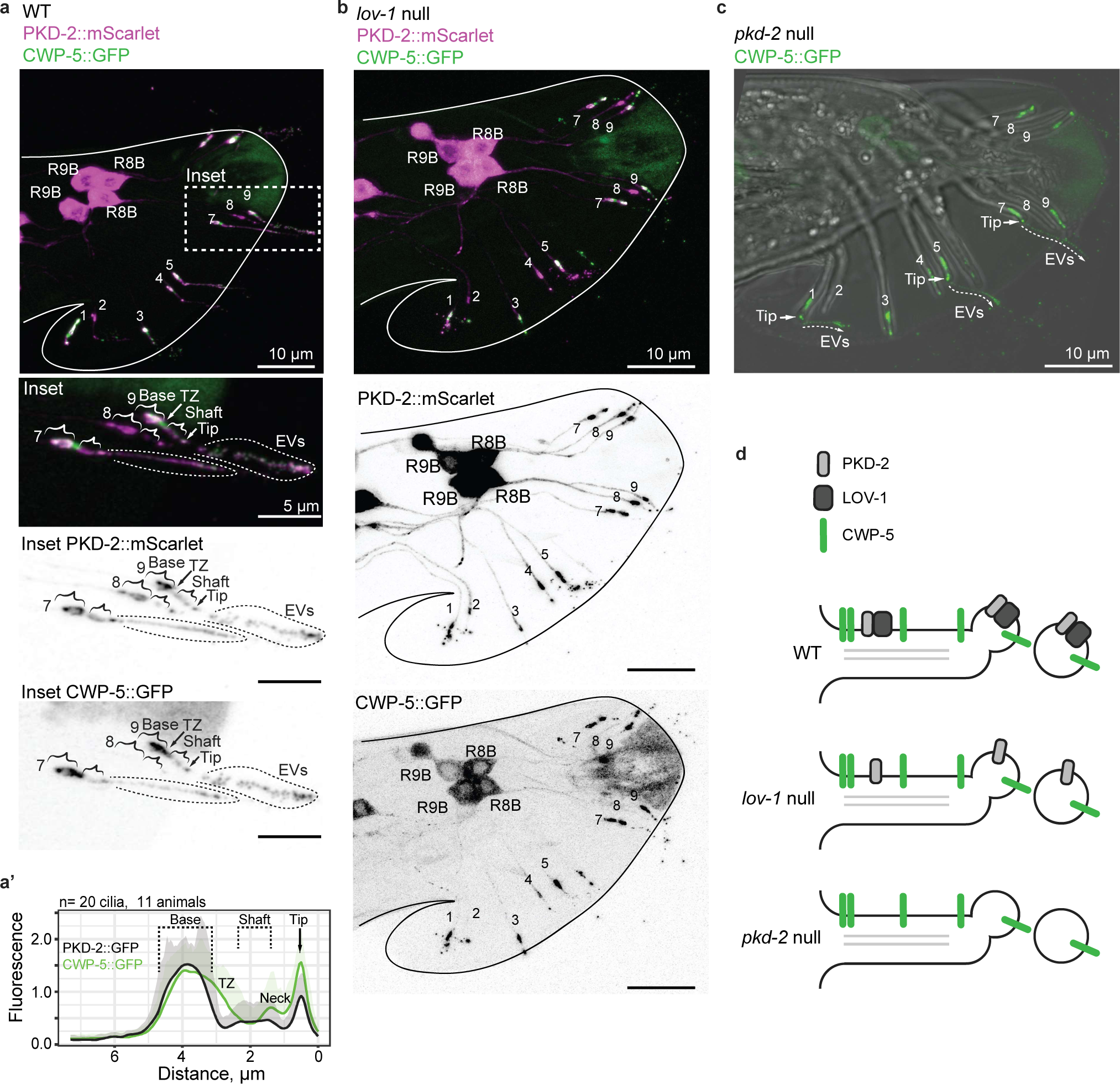
The polycystin complex is not required for shedding of ciliary EVs as evidenced by tracking cargo CWP-5. **a,** Flattened z-stacks showing colocalization of PKD-2::mScarlet with CWP-5::GFP with a zoomed-in inset. **a’,** Representative average fluorescent profile of CWP-5::GFP and PKD-2::mScarlet along the cilium. Note the prominent presence of CWP-5::GFP in the areas depleted of PKD-2::mScarlet – in the transition zone (TZ) and the neck of the cilium, **b-c,** Flattened z-stacks showing that disruption of *lov-1* (**b**) and *pkd-2* (**c**) does not alter CWP-5::GFP ciliary and EV localization. **d,** Scheme of molecular mechanism showing that loading CWP-5::GFP to cilia and ciliary EVs is independent of the polycystins.

Unlike PACL-1 and PAMLs, CWP-5 did not exhibit complete colocalization with PKD-2 within the cilium. Specifically, CWP-5::GFP was enriched in the transition zone and the neck compartment proximal to the ciliary tip, two exclusion areas for PKD-2 reporters (Fig. 6a-a’, Extended Data Fig. 9b). The ciliary localization of CWP-5::GFP was independent of LOV-1 and PKD-2 (Fig. 6b-c), indicating that CWP-5::GFP is transported to the cilium and loaded to polycystin-carrying EVs independently from the polycystin complex (Fig. 6d).

## Discussion

In this study, we used TurboID labeling to discover cargo associated with PKD-2 within ciliary EVs of *C. elegans*. We took a modular approach generating TurboID with an anti-GFP nanobody domain tethering the proximity labeling enzyme to a GFP-labeled cargo of interest. This EV-TurboID approach has many implications for the future of the EV field as opposed to direct fusions. Expression of the nanobody::TurboID under cell– or tissue-specific promoters in strains with endogenously GFP-tagged proteins enabled us to address EV cargo composition in a cell-specific manner. Our previous work shows that EVs with shared cargo can be generated via distinct routes and thus might differ drastically in content ^16^. The application of modular proximity labeling in cell– and tissue-specific manner is a robust strategy for dissecting the composition of EV subtypes and individual EVs with unprecedented resolution.

Our work shows the regulated loading of cargo onto ciliary EVs at a single-cell, single-EV level in *C. elegans*. We discovered a hierarchical assembly of signaling components on ciliary EVs, demonstrating how LOV-1 recruits partners and cargo to cilia and ciliary EVs. An association with LOV-1 was essential and a pre-requisite for EV cargo loading. LOV-1 assembled with transmembrane partners (PACL-1, PAML-1, PAML-2) in the cell body and TRAFs TRF-1 and TRF-2 within the ciliary shaft. Conversely, CWP-5 ciliary and EV localization was polycystin-independent and via an unidentified mechanism. This level of precision in analyzing individual EV components and EV cargo loading *in vivo* remains unattainable in vertebrate models so far.

We propose a groundbreaking concept: a single genetic change (like a polycystin-1 mutation) significantly alters EV content and potential signaling properties (Figure 7). EVs have been implicated in the pathogenesis of ADPKD^29^. Clinical observations of polycystin-1 mutations causing more severe polycystic kidney disease than the polycystin-2 patients^30^ align with our findings, suggesting distinct EV content in different polycystin mutation patients, potentially impacting disease progression. Intriguingly, soluble fragments of the PC1 C-type lectin ectodomain act as a ligand and activate the polycystin complex in cultured mammalian cells^31^, suggesting that polycystins may signal between cells using EVs. Ward and colleagues proposed using human urinary EVs as a source for proteins that interact with the polycystin complex^32^.

**Fig. 7.**
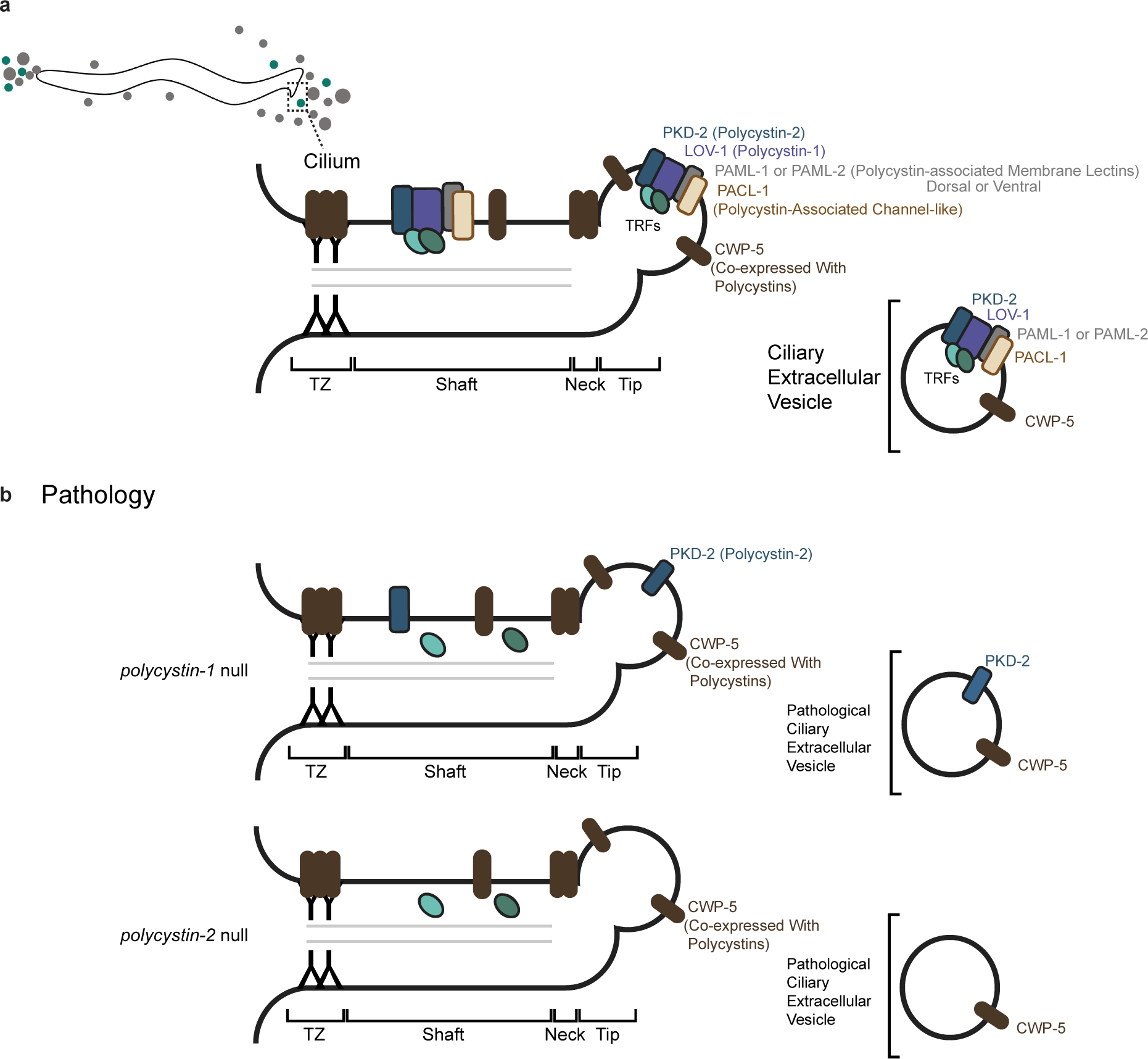
Summary of cargo loading to polycystin ciliary EVs of *C. elegans*. A single genetic perturbation leads to a drastic change in EV cargo composition. Compare WT polycystin EVs (**a**) with EVs of the *lov-1* and *pkd-2* mutants (**b**).

Our work highlights the importance of understanding polycystin function in EVs and the potential application of single EV analysis for non-invasive early diagnosis of ADPKD.

To our knowledge, this work is the first evidence of the physical presence of TRAF homologs (TRFs) within the cilium. Specifically, TRF-1 (an adaptor with a ubiquitin-ligase-like RING domain) is required for cytoplasmic abundance of TRF-2 (a TRAF without RING) and its entrance to the ciliary shaft. In contrast, TRF-1 can enter the cilium independently of TRF-2 but depends on it for its mutual loading to polycystin EVs. Neither TRF-1 nor TRF-2 is required to release the polycystin complex in the form of ciliary EVs, supporting the hypothesis that cargo loading into ciliary EVs is hierarchical.

Given the high conservation of the TRAF family and the polycystins between humans and *C. elegans*, their functional interactions might likewise be conserved. This hypothesis is supported by the fact that *Traf6* knock-out mice phenocopy skeletal ciliopathies, including exencephaly, problems with neural tube closure, shortening of long bones and tooth agenesis in neonates, focal alopecia, delayed pigmentation, and underdevelopment of skin glands, including sweat and sebaceous glands ^33–36^. These phenotypes mimic ciliopathies of human, such as *NEK1*, *DYNC2H1*, and *IHH* mutations (shortening of long bones), tooth agenesis in patients with BBSome variants, sparse hair and focal alopecia of *EDA* patients ^37–42^. Our work and the literature suggest that human TRAFs might act in the cilium, possibly in tandem with polycystins. Consistent with this idea, TRAF7 mutation causes skull-base meningiomas and congenital heart disease in humans, and is important for intraflagellar transport and ciliogenesis in *Xenopus*^43^.

Additionally, our study establishes the link between the polycystins and a channel-like PACL-1. Polycystin dysfunction in ADPKD patients is associated with deregulated monovalent ion transport that directly contributes to cyst enlargement ^44^. This association suggests that disrupted polycystins in human might cause mislocalization of ion channels, resulting in deregulated ion homeostasis and abnormal fluid accumulation.

Polycystins’ connection with transmembrane C-type lectins of dorso-ventral specialization hints at conserved involvement of polycystins in the sensation of spatial orientation as observed in cerebrospinal fluid-contacting neurons (CSF-cNs) of zebrafish ^3–5^, and in the left-right organizer of the early mouse embryo ^6^. In *C. elegans*, polycystin-associated PAMLs’ might signal the male during mating regarding the vulva’s location based on their differential affinities to hermaphrodite cuticle glycans along the body^45^. This aspect of polycystin biology warrants further investigation.

In summary, our study presents a new application of proximity labeling for EV subtyping that we used to precipitate *bona fide* cargo of polycystin EVs of *C. elegans*. We established a connection between polycystins and three classes of proteins (channel, C-type lectin transmembrane, and TRAFs). Our study offers a blueprint for further exploration of EV biology.

### Limitations of study

Teasing out the biological activity of EV subtypes and determining the functions of individual EV cargo components are extremely challenging but critical questions to tackle. We previously demonstrated that a heterogenous mixture of isolated EVs triggers changes in male locomotory tail-chasing behavior and that EVs isolated from a cargo sorting/shedding mutant fail to elicit a behavioral response. We are currently developing methods to specifically isolate polycystin-carrying EV subtypes, which can then be genetically depleted for individual cargo components and assayed for bioactivity. We are also focused on identifying the EV target cells and specific bioactivities.

## Acknowledgements

We are grateful to Haiyan Zheng and David Sleat for assistance with the mass spectrometry analysis; Gloria Androwski and Helen Ushakov for outstanding technical assistance; Premal Shah and John Favate for sharing and maintenance of the BioComp fractionator that was instrumental in this work, Laura Bianchi, Paul DeCaen, and Monica Driscoll for discussions; members of the Rutgers *C. elegans* community for thought-provoking questions. We also thank WormBase^46^ and WormBook which were used daily during this project. The work was funded by grants from the National Institutes of Health (NIH) DK116606, DK059418, and NS120745 to M.M.B.; NIH K12 GM093854 INSPIRE (IRACDA NJ/NY) postdoctoral fellowships to K.C.J, and intramural pilot Rutgers Core Facility grant to I.A.N. Protein identification was performed in the Biological Mass Spectrometry Facility of Robert Wood Johnson Medical School at Rutgers University supported by the NIH instrumentation grants S10OD025140 and S10OD016400 with intramural grants to I.A.N. Some strains were provided by the Caenorhabditis Genetics Center (CGC), which is funded by NIH Office of Research Infrastructure Programs (P40 OD010440). The Center for *C. elegans* Anatomy and WormAtlas provided valuable anatomical and ultrastructural resources (R24OD010943).

## Author Contributions

Conceptualization, I.A.N. and M.M.B.; methodology, I.A.N., J.W., J.D.W, M.M.B.; investigation, I.A.N., E.D., K.C.J, J.S.; visualization, I.A.N..; funding acquisition, M.M.B., I.A.N.; project administration, I.A.N., M.M.B.; supervision, I.A.N., J.W., M.M.B.; writing – original draft, I.A.N.; writing – review & editing, I.A.N., K.J.C., J.W., and M.M.B.

## Figure Titles and Legends

**Extended Data Fig. 1.**
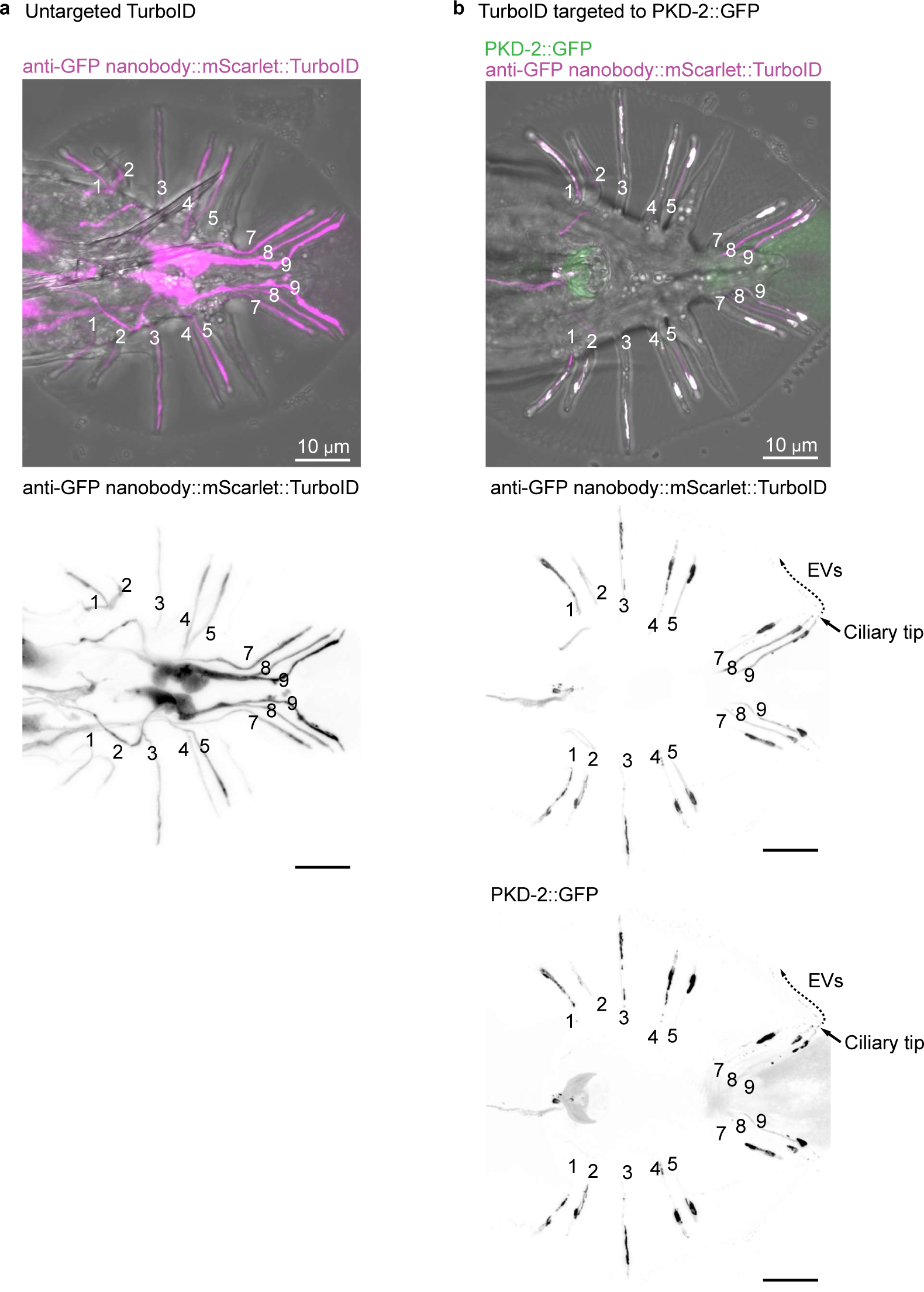
Targeting TurboID module to PKD-2::GFP (related to Fig. 1). **a,** anti-GFP nanobody::mScarlet::TurboID is uniformly abundant in cell bodies, neurites, and cilia in an untargeted state in the absence of any GFP-labeled protein. **b,** Coexpression of the *anti-GFP nanobody::mScarlet::TurboID* and *pkd-2::GFP* results in specific targeting of the TurboID module to PKD-2::GFP subcellular locations, including cilia and ciliary EVs. Male tails in panels **a** and **b** are oriented in different positions (dorsal on **a**, ventral on **b**), so cell bodies are not captured in panel **b**.

**Extended Data Fig. 2.**
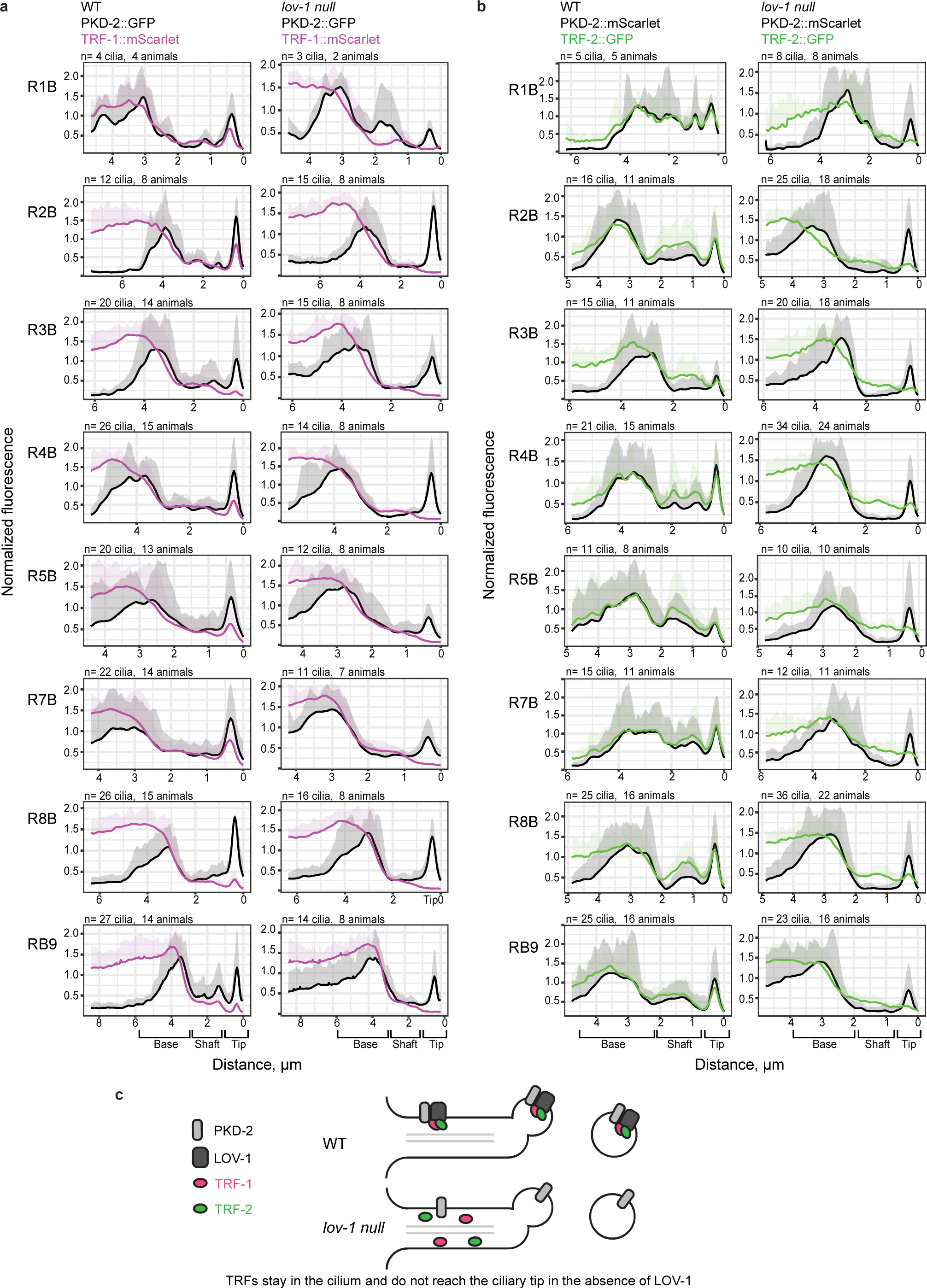
LOV-1 is required for loading TRFs to ciliary PKD-2 EVs (related to Fig. 2). **a-b,** Average fluorescence profiles through cilia of RnB neurons of WT and *lov-1* null males for the pair of PKD-2::GFP and TRF-1::mScarlet (**a**) and for PKD-2::mScarlet and TRF-2::GFP (**b**). **c,** Scheme of molecular mechanism loading TRFs to polycystin EVs. Summary of findings.

**Extended Data Fig. 3.**
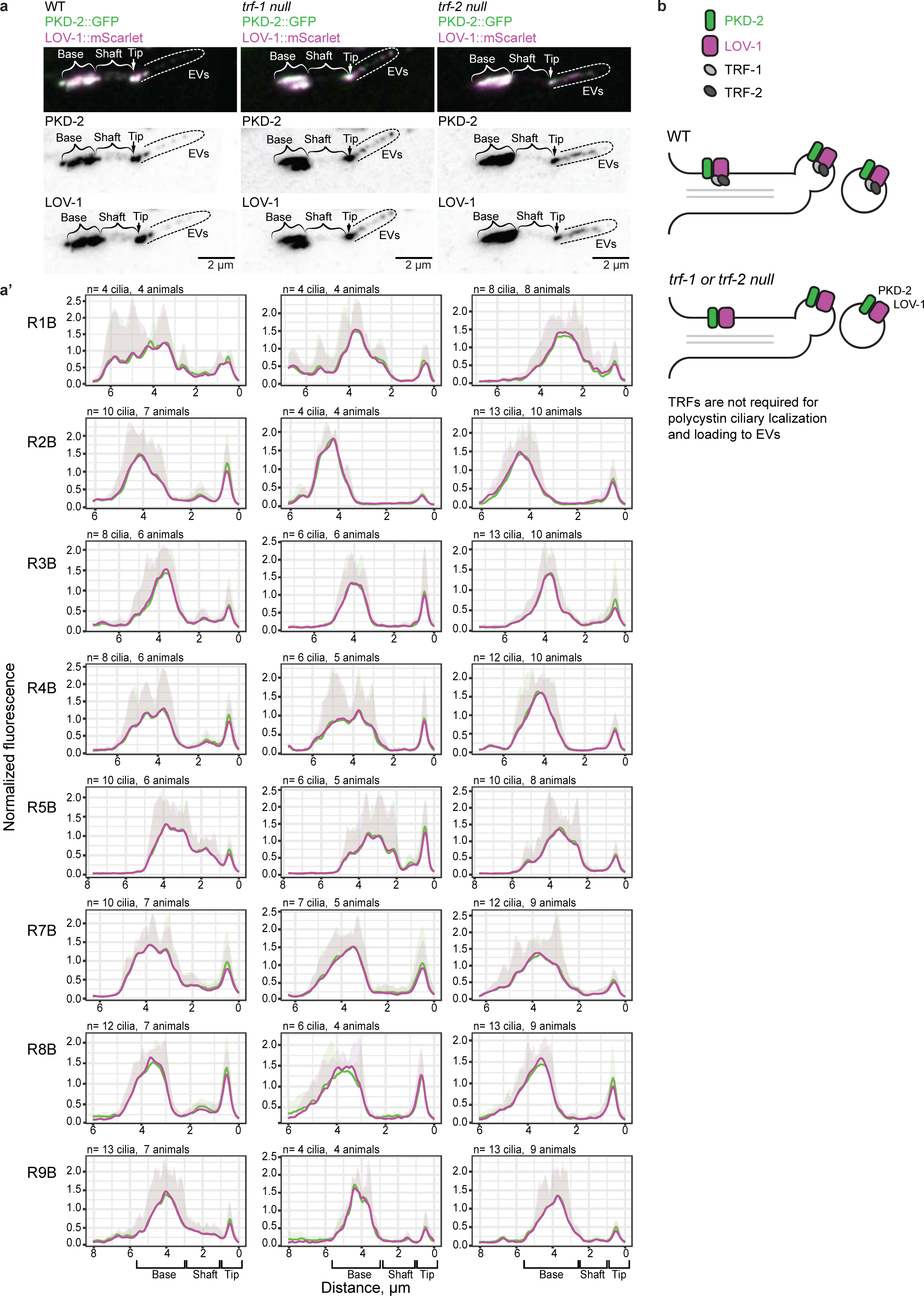
TRFs are not required for polycystin EV release. **a,** Flattened z-stacks showing colocalization of PKD-2::GFP and LOV-1::mScarlet in the cilium and on EVs in WT, *trf-1* null, and *trf-2* null animals. **a’,** Average fluorescence profiles through cilia of RnB neurons. **b,** Summary of findings.

**Extended Data Fig. 4.**
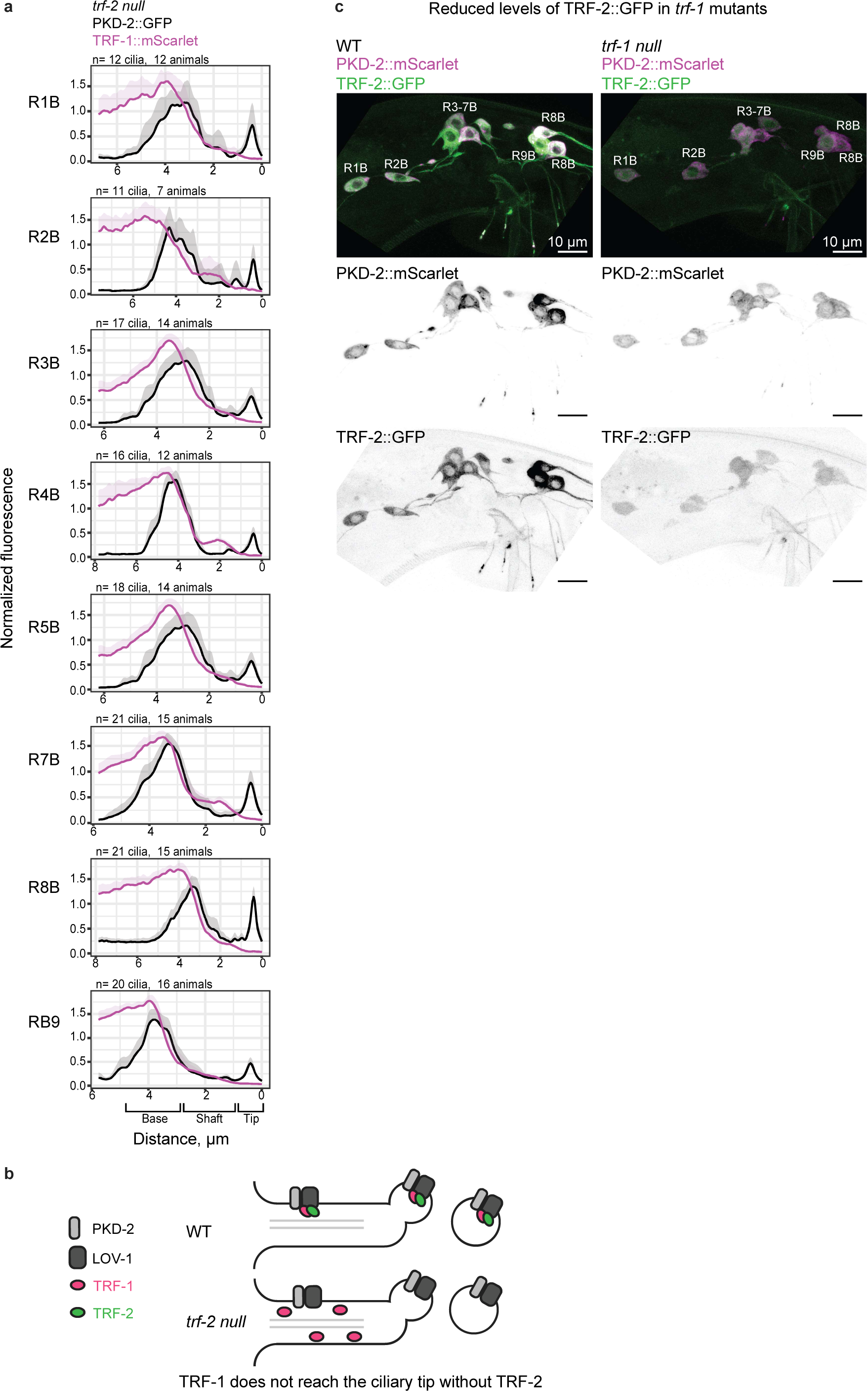
TRFs require each other for their loading to ciliary EVs. **a,** Flattened z-stacks showing average fluorescence profiles of PKD-2::GFP and TRF-1::mScarlet along the cilium of *trf-2* null animals. TRF-1 is depleted form the ciliary tip and is not loaded to PKD-2::GFP EVs. **b,** Summary of findings. **c,** Flattened z-stacks (uniformly adjusted) showing representative images of TRF-2::GFP and PKD-2::mScarlet levels in the cell bodies of RnB neurons. In the *trf-1* null mutant, TRF-2::GFP levels are reduced three times.

**Extended Data Fig. 5.**
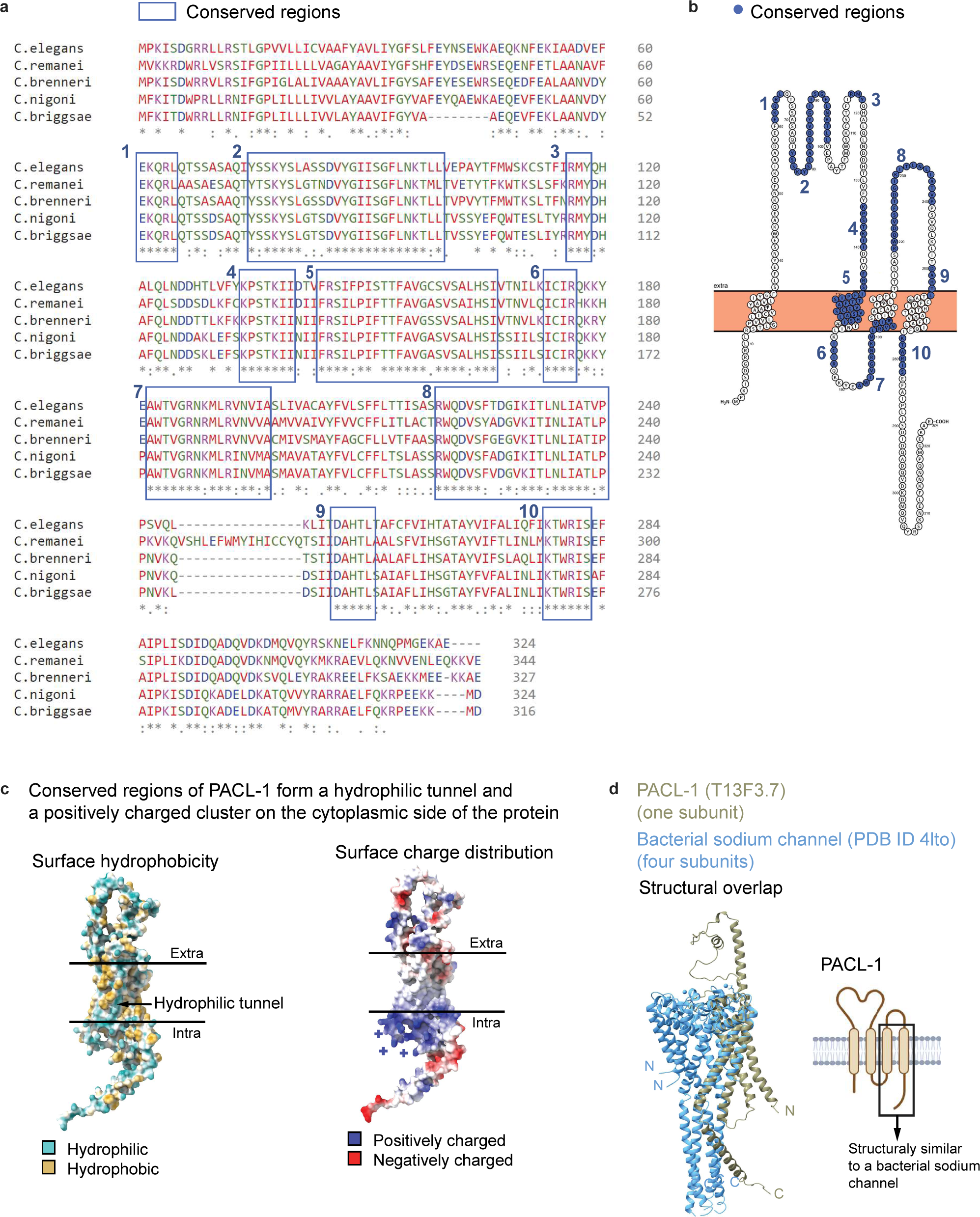
PACL-1 predicted 3D structure exhibits channel-like properties. **a**, Alignment of PCAL-1 homologs within the *Caenorhabditis* genus. Regions of the high similarity are boxed and numbered. **b,** Diagram of PACL-1 amino acid sequence, regions of the highest similarity from panel **a** are shown as blue-shaded amino acid stretches. **c,** Surface hydrophobicity of PACL-1 (arrow points to the predicted hydrophilic tunnel) and charge distribution (positively-charged cluster on the cytoplasmic part is in blue).**d,** Structure overlay of the bacterial sodium channel (four subunits, blue) and PACL-1 AlphaFold predicted structure (a single subunit, yellow). Our search in the Protein Data Bank for structural folds akin to the AlphaFold-predicted structural model of PACL-1 yielded a bacterial sodium channel (PDB ID: 4lto) as a candidate (**a, c**). The structural similarity of PACL-1 and the bacterial sodium channel is confined to transmembrane helices 3 and 4, connected by an extracellular loop (**a, c**); all are regions less conserved among *Caenorhabditis* species (**b, c**). These helices potentially might stabilize the conserved amphipathic helix. Thus, specific amino acid composition within these regions may be less critical for protein function. Examination of the evolutionary conservation of PACL-1 across *Caenorhabditis* species revealed almost invariable conservation within specific segments, notably an amphipathic helix (a putative pore – region 5 on panels **a** and **b** forms hydrophilic tunnel on panel **c**) and a positively charged cluster located on the cytoplasmic side of the predicted pore entrance (region 7 and 10 on panels **a** and **b**, blue cluster on panel **c**) that might serve as an “anion sink.” Among the sequence homologs of PACL-1 outside of the *Caenorhabditis* genus were a four-pass transmembrane protein A0A016SLD6_9BILA (*Acey_s0209.g2096* gene) of *Ancylostoma ceylanicum* (intestinal hookworm) that distantly resembles TWiK potassium channels (Tandem of P-domains in a Weakly Inward rectifying K^+^ channel). Taken together, these observations suggest a role for PACL-1 as a channel. Whether PACL-1 functions as an anion or cation channel necessitates experimental validation beyond the scope of this study.

**Extended Data Fig. 6.**
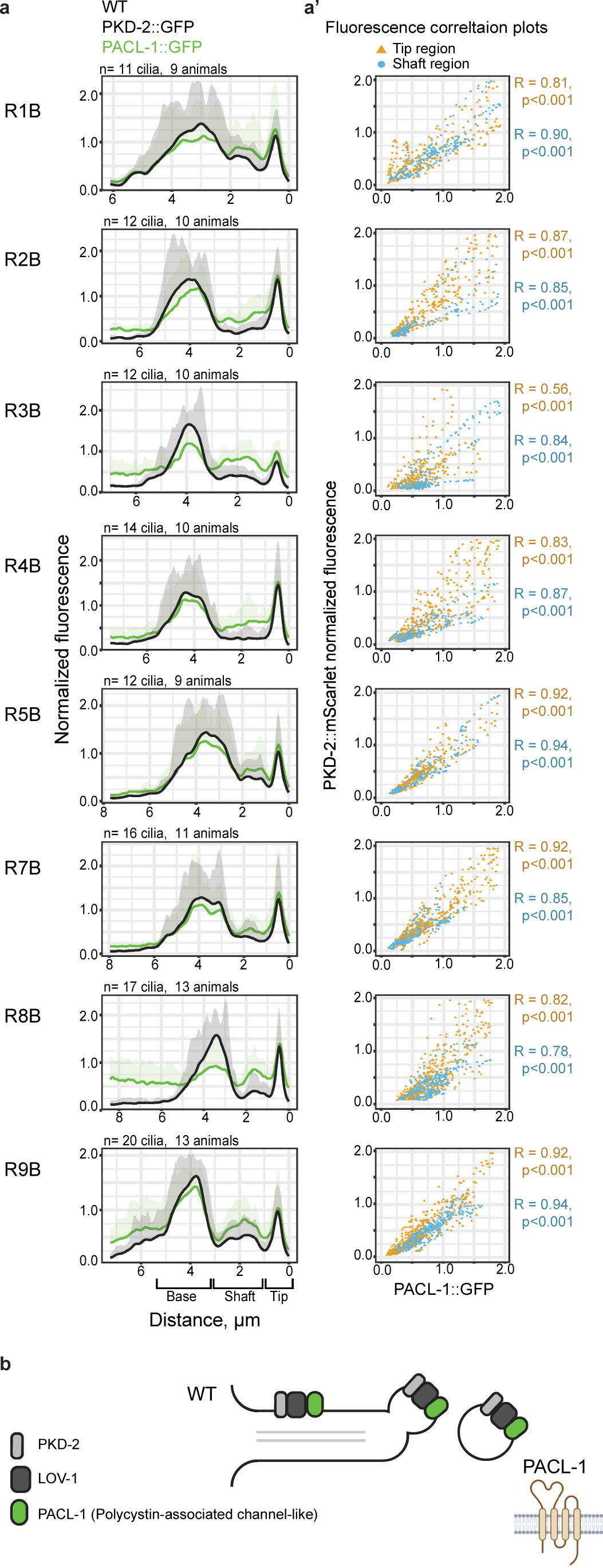
PACL-1::GFP colocalizes with PKD-2::mScarlet in cilia and EVs. **a,** Fluorescence profiles through cilia of RnB neurons. **a’,** Colocalization plots show a high level of correlation between the PACL-1:GFP and PKD-2::mScarlet fluorescence intensities in the ciliary shaft (blue) and ciliary tip (yellow). **b,** Summary of molecular mechanism. PACL-1 associates with the polycystin complex and is released in EVs.

**Extended Data Fig. 7.**
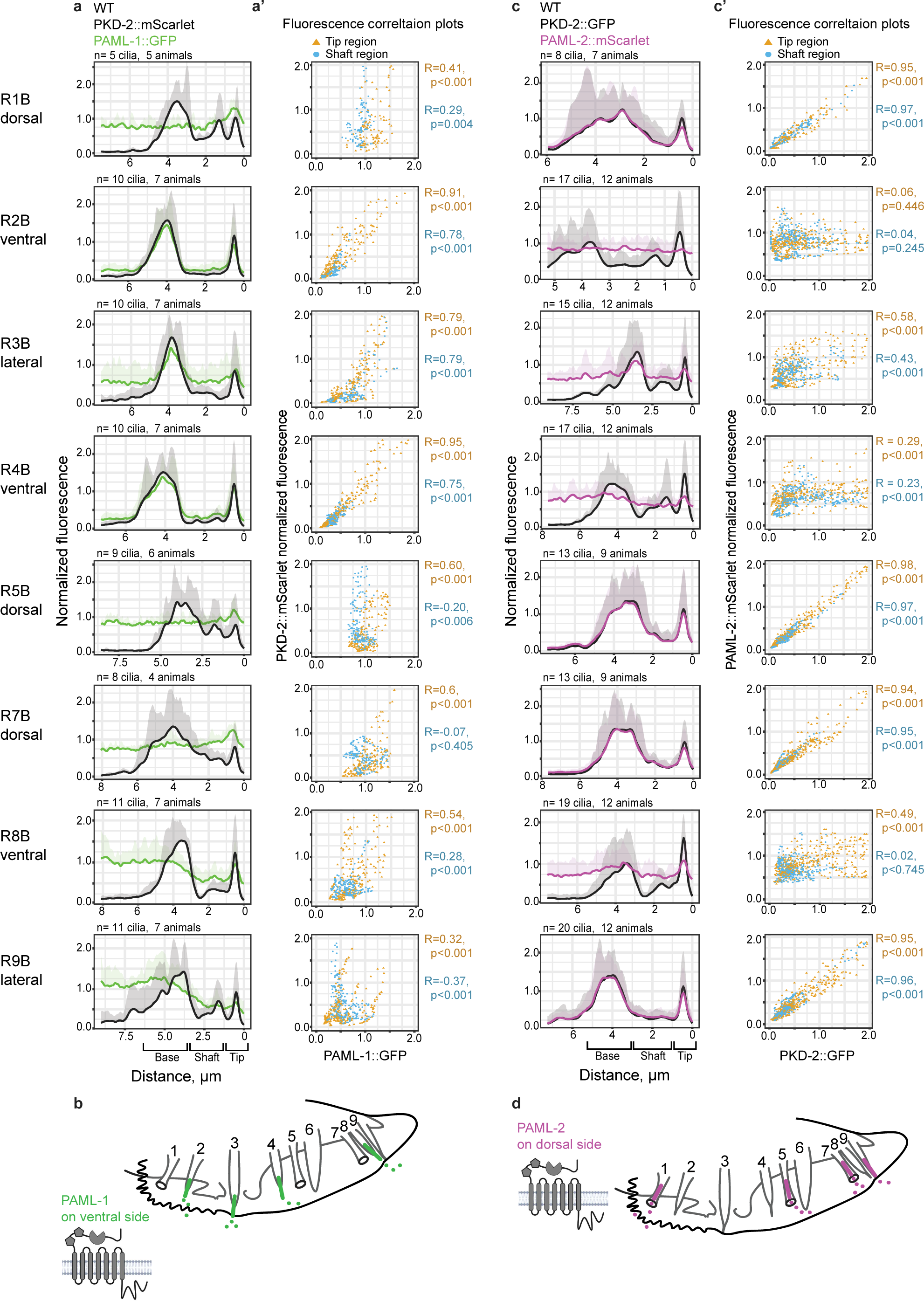
PAML-1 and PAML-2 are polycystin-interacting transmembrane C-type lectins with dorso-ventral specialization. **a**, Fluorescence profiles through cilia of RnB neurons for PAML-1::GFP. **a’,** Colocalization plots showing a correlation between the PAML-1::GFP and PKD-2::mScarlet fluorescence intensities in ciliary shaft (blue) and ciliary tip (yellow). **b,** Summary cartoon for PAML-1 presence in cilia and EVs. **c,** Fluorescence profiles through cilia of RnB neurons for PAML-2::mScarlet. **c’,** Colocalization plots showing a correlation between the PAML-2::mScarlet and PKD-2::GFP fluorescence intensities in ciliary shaft (blue) and ciliary tip (yellow). **d,** Summary cartoon for PAML-2 presence in cilia and EVs. Fluorescence values are normalized to the average of minimum and maximum values for each cilium.

**Extended Data Fig. 8.**
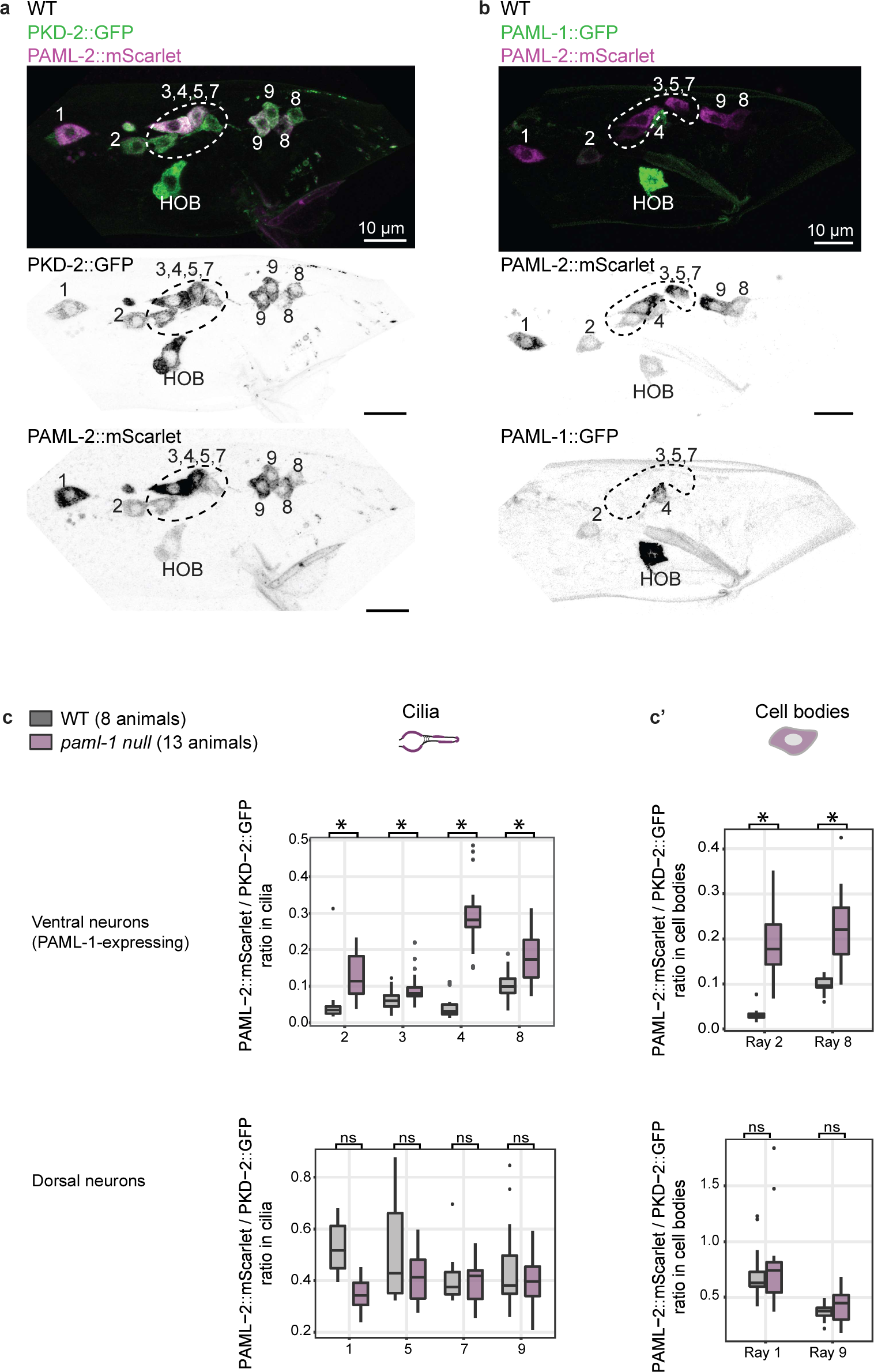
Cell-specific expression of PAML-1 in ventral rays 2, 3, 4, and 8 inhibits expression of PAML-2, resulting in PAML-2 ciliary localization only in dorsal rays 1, 5, 7, and 9. **a,** All PKD-2::GFP-expressing neurons express PAML-2::mScarlet. Image taken at the L4 molt stage (the last molting stage at which most genes required for sexual maturity start their expression). **b,** Expression patterns of PAML-1::GFP and PAML-2::mScarlet at the L4 molt stage. While PAML-2 is present in all neurons, PAML-1 expression is restricted to a ventral subset of the neurons. At this stage, ray 2, 4, and HOB cell bodies are PAML-1-positive. **c,** The *paml-1* null mutation results in ectopic ciliary localization of PAML-2 in ventral rays 2, 3, 4, 8 (upper left panel). The ciliary presence coincides with increased paml-2::mScarlet presence in cell bodies of those neurons (upper right panel). We measured the cell bodies of neurons 2 and 8 because they are easily identifiable by location; the proximity of neuronal cell bodies 2 and 4 to neurons 5 and 7 made their identification ambiguous, and thus, we did not take these measurements. In dorsal neurons (lower panels), levels of PAML-2 in cilia and cell bodies were unaffected by the *paml-1* null mutation. *Mann-Whitney U Test *p* < .00001 We propose that the mechanism of selective transport of PAML-1 to the cilium of ventral neurons stems from the different affinities of PAML-1 and PAML-2 for the polycystin complex. In ventral neurons that express both *paml-1* and *paml-2*, only PAML-1 is recruited to the cilium, presumably due to its strong binding to the polycystin complex. Conversely, PAML-2 likely has lower affinity to polycystins and thus binds the polycystin complex solely in the absence of PAML-1, as observed in dorsal neurons. This mechanism produces two distinct types of polycystin-carrying ciliary EVs, originating from dorsal and ventral neurons, carrying either PAML-1 or PAML-2 cell-specific markers.

**Extended Data Fig. 9.**
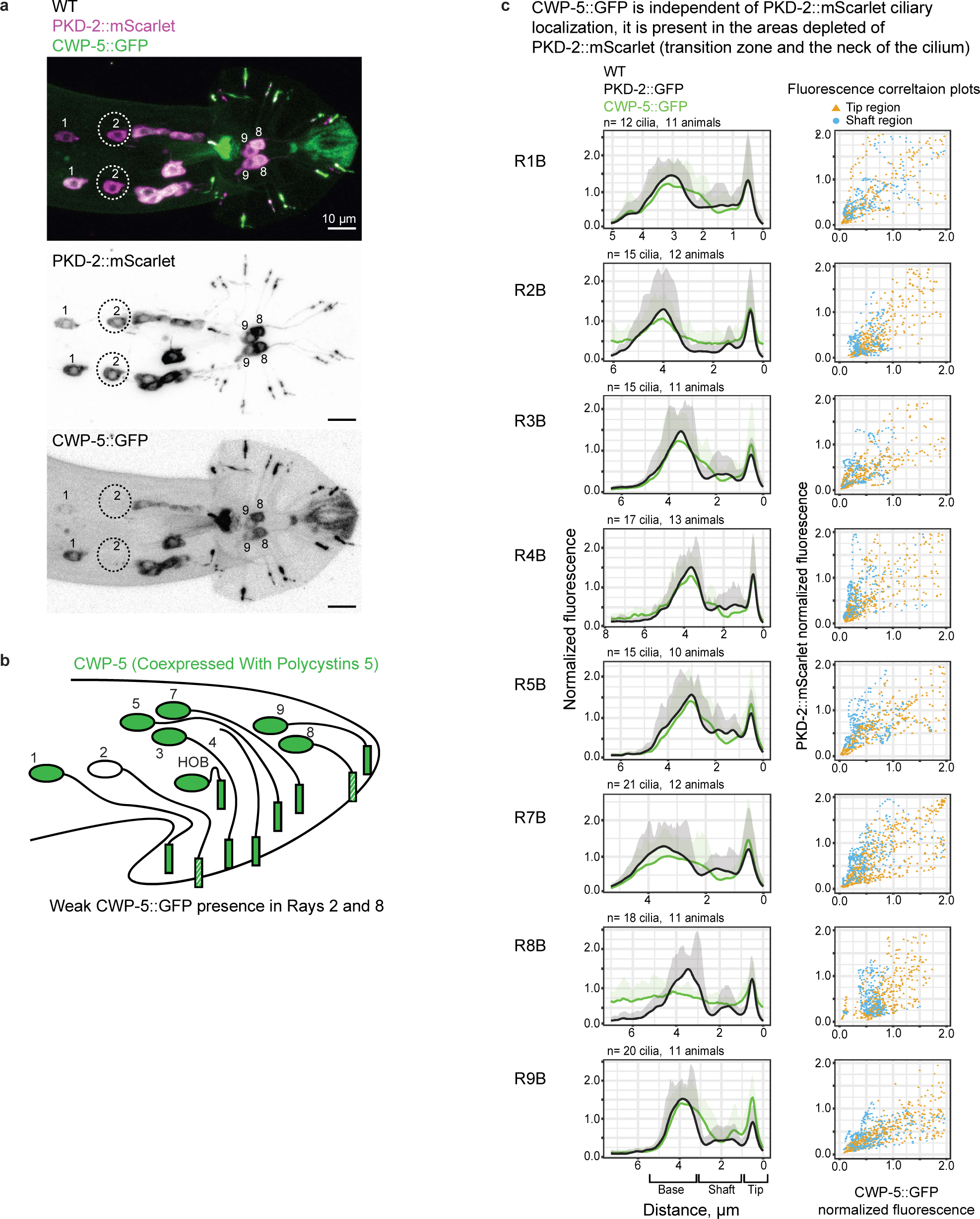
CWP-5 expression pattern in PKD-2 expressing neurons displays ray-specificity. **a,** Flattened z-stack showing PKD-2::mScarlet and CWP-5::GFP expression. Note the absence of CWP-5::GFP in the cell body of ray 2. **b,** CWP-5::GFP summarized expression pattern. c, Normalized fluorescence profiles through cilia of RnB neurons for CWP-5::GFP and PKD-2::mScarlet. Note the shouldering effect of CWP::GFP profile compared to PKD-2::mScarlet in the transition zone and the area proximal to the ciliary tip. **c’,** Colocalization plots show less correlation between PKD-2::mScaret and CWP-5::GFP compared to PACL-1::GFP (Extended Data Fig.5a’), suggesting that CWP-5 has ciliary functions independent of the polycystins.

**Extended Data Fig. 10.**
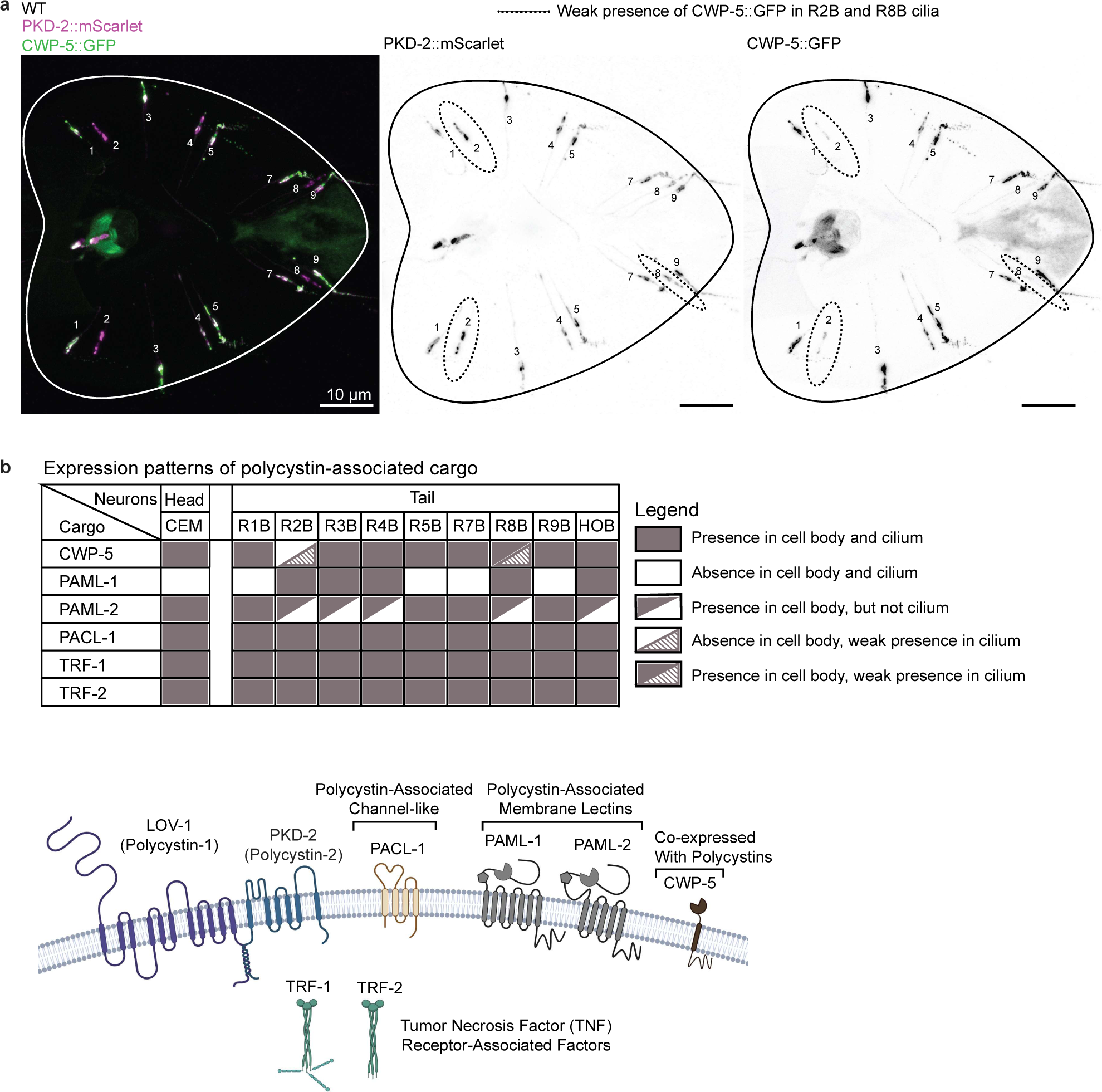
CWP-5::GFP ciliary localization in ray 2 and 8 is observed in 50% cases. **a,** Representative image showing CWP-5::GFP presence in cilia or R2B and R8B neurons (circled). **b,** Summary of PKD-2 interactor expression patterns. Within the R2B and R8B neurons, CWP-5::GFP showed weak and inconsistent ciliary presence (observed at low levels in 50% cases: 19 out of 26 R2B cilia were CWP-5::GFP positive despite no detectable fluorescence in the R2B cell body, and 28 out of 50 R8B cilia were weakly CWP-5::GFP-positive despite high levels of CWP-5::GFP in the R8B cell body). These results suggest the existence of molecular mechanisms inhibiting transport of CWP-5 from the R8B cell body to the cilium. The source of CWP-5::GFP in the R2B cilium was unclear because CWP-5::GFP was not present in the R2B cell body. We propose three explanations for the absence of CWP-5 in the cell body and its presence in the cilium of the R2B neuron. First, R2B might express CWP-5 at exceedingly low levels, evading detection by our imaging technique. However, CWP-5 relocation to the cilium might be so efficient that even minute amounts in the cell body would lead to detectable accumulation within the cilium. Second, synthesis of the CWP-5 protein might occur exclusively at the ciliary base of R2B, resulting in its exclusive presence in the cilium and not the cell body. Third, the accumulation of ciliary CWP-5 in R2B cilia might arise from the fusion of CWP-5-carrying environmental EVs with the cilium of the R2B neuron. The latter scenario suggests that the R2B neuron might function as a recipient cell for EVs released from other rays of the same or different animal.

## Tables with titles and legends

Extended data Table 1.xls file with mass spectrometry results of protein identification followed by the pulldown of the biotin-labeled EV cargo.

## Extended data – Materials and Methods

### Strains and culture conditions

Animals were cultured at 20°C using the OP50 *E.-coli* strain as a food source. For culturing animals for EV isolation, see sections below.

**Table S2.**
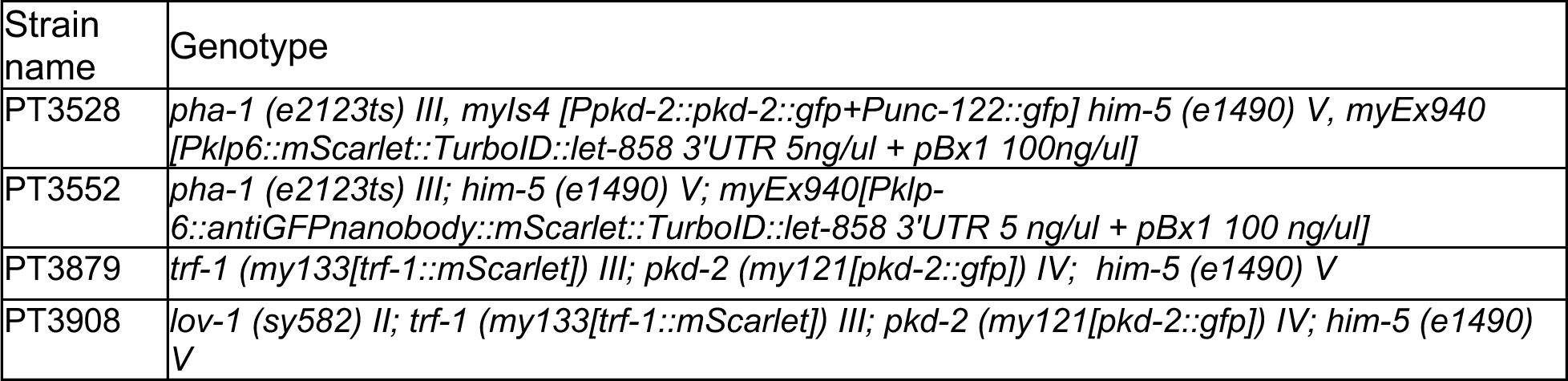
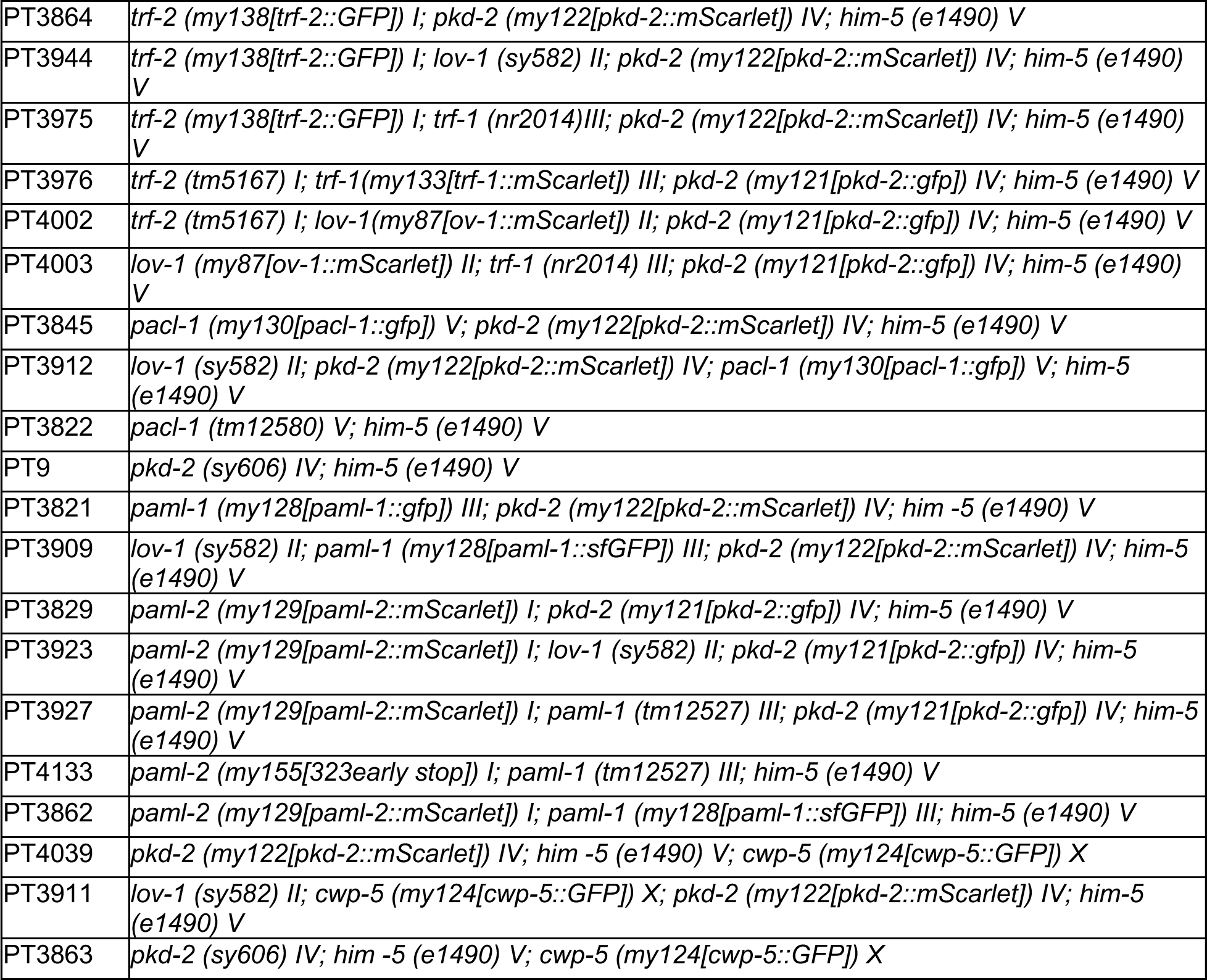
List of strains used in the study.

### TurboID-based enzymatic labeling of neuronal ciliary EVs in *C. elegans*

The overall scheme of the experiment included the isolation of EVs from two strains. The control strain carried TurboID under the *klp-6* promoter driving expression in IL2, CEMs, RnBs, and HOB neurons. Thus, in the control strain, TurboID was not targeted at any specific cell location, its localization looked uniform throughout the neurons, including cell bodies, nuclei, axons, dendrites, and cilia (Extended Data Fig. 1a). The sample strain carried TurboID under the *klp-6* promoter and the *Ppkd-2::pkd-2::gfp* transgene expressed in the *pkd-2*-specific CEMs, RnBs, and HOB neurons. In this sample strain, TurboID followed the subcellular localization of PKD-2::GFP in cell bodies, cilia, and was specifically loaded and enriched in PKD-2::GFP carrying EVs (Fig. 1c, Extended Data Fig. 1b).

To initiate the culture, 240 hermaphrodites at the larval stage 4 were transferred to 40 6-cm normal growth medium (NGM) plates (6 animals per plate) and grown for two generations. On day 6, each plate was chunked into quarters. The chunks were transferred into 15-cm plates with high growth medium (HGM) (3 g of NaCl, 2.5 g of peptone, and 20 g of agar per 1 L supplemented with 4 mL of cholesterol stock (5 mg/mL of ethanol), 1 mL 1 M CaCl_2_, 1 mL 1 M MgSO_4_, and 25 mL 1M potassium phosphate buffer pH 6.0) seeded with a full lawn of E. coli OP50 (1.8 mL/plate). Animals were allowed to propagate for two more generations at 22°C until they consumed most of the bacteria and yet were not starved. In total, about 4 million of adult worms were used to isolate EVs for each replicate.

### Harvesting EVs of *C. elegans*

On the day of EV harvest, we washed *C. elegans* off the HGM plates with the M9 buffer (3 g KH_2_PO_4_, 6 g Na_2_HPO_4_, 5 g NaCl, 1 mL of 1 M MgSO_4_ per 1 L) into 15 mL conical tubes. We centrifuged the collected animal suspension at 3,000 g for 15 min to pellet the worms and collect the EV-containing supernatant. To increase EV yields, we resuspended animals in fresh M9 buffer and repeated the centrifugation step (15 min at 3,000 g) twice, collecting the supernatant with released EVs each time.

To clear the EV-containing supernatant of residual bacteria and large debris, we centrifuged it at 10,000g (8,000 rpm in SW28 swinging bucket rotor, Beckman Coulter) for 30 minutes at 4°C. We repeated this step 3 times, collecting the more clarified supernatant. After the final clarification step, we pelleted EVs from the cleared supernatant onto the 36% iodixanol cushion at 100,000g (28,000 rpm in SW28 swinging bucket rotor, Beckman Coulter) for 2 hours at 4oC. The cushion was prepared by mixing three parts of 60% iodixanol (OptiPrep, Sigma #D1556) and two parts of 8% sucrose to achieve the final density of 1.2 g/L.

To concentrate pelleted EVs into a smaller volume for further loading onto density gradients, we resuspended the collected EVs in the M9 buffer. We pelleted the EV suspension again in a smaller volume tube at 100,000g (30,000 rpm SW41 swinging bucket rotor, Beckman Coulter) onto the 36% iodixanol cushion.

Before loading the concentrated EV suspension onto density gradients, we ensured the density of the suspension was less than 1.1 g/mL by diluting it with M9 buffer (typically, that meant adding 2 mL of fresh M9 buffer to 1 mL of concentrated EV suspension). Using a thin glass Pasteur pipette, we loaded the diluted EV suspension onto a top of density gradients (1.1 –1.2 g/mL), prepared by a freeze-thaw method described in Nikonorova *et al.* 2022 ^16^. To drive TurboID-labeled EVs to their respected density within the tube, we conducted isopycnic centrifugation at 100,000g (30,000 rpm, 16 hours, at 4°C).

To collect TurboID-containing EVs, we fractionated the equilibrated gradients on the BioComp Piston Gradient Fractionator into 30 fractions of 375 mL each. To examine fractions for PKD-2::GFP and anti-GFP nanobody::mScarlet::TurboID EVs using a super-resolution microscope (Zeiss LSM880 with Airyscan). We layered an array of representative droplets from each fraction onto a glass slide. We used double-adhesive stickers to form chambers for the droplets (SecureSeal spacers by Electron Microscopy Sciences, Cat# 70327-9S). Once the most enriched fractions were identified, we proceeded to the extraction of biotinylated proteins.

### Extraction of biotinylated proteins from *C. elegans* EVs

We closely followed the protocol of Branon *et al*.^17^. Specifically, EVs were lysed with RIPA buffer and then incubated with streptavidin-conjugated beads (Pierce Streptavidin Magnetic beads, ThermoFisher #88816; 25 μl of the bead suspension per 300 μg of total protein). Afterward, beads were washed twice with the RIPA buffer, once with 1 M KCl, once with 0.1 M Na_2_CO_3_, once with 2 M urea in 10 mM Tris-HCl (pH 8.0), and five times with 50 mM NH_4_HCO_3_. We transferred the beads to a fresh tube during the final wash. The beads were then submitted to the Rutgers Proteomics facility for on-bead trypsin digestion and protein identification.

### Generation of endogenously labeled strains

To validate mass spectrometry findings, we generated endogenous reporters using CRISPR/Cas9-mediated genome editing as described in Dokshin et al. 2018^47^. All generated strains were verified by Sanger sequencing and are reported below.

### Super-resolution fluorescence imaging and analysis

For *in vivo* imaging, we anesthetized *C. elegans* males with 10 mM levamisole solution in the M9 buffer and mounted animals onto thin 10% agarose pads. Animals were imaged within 30 minutes of their mounting. Super-resolution imaging was performed on a Zeiss LSM880 confocal system equipped with Airyscan Super-Resolution Detector, Single Photon Lasers (488 nm for GFP and 561 nm for mScarlet), motorized XY Stage with Z-Piezo and T-PMT. For analysis of fluorescence profiles through cilia of RnB neurons, fluorescence values were normalized to the average of minimum and maximum values for each cilium.

### Assessment of male mating behavior

For mating assays, we used fresh OP50 lawns of 10 μl each, seeded 2 hours before the assay, dried at 37°C incubator, and prechilled to the room temperature. We then populated the mating spots with ten 1-day old hermaphrodites bearing *unc-31* null mutation that reduces movement, making hermaphrodites easy mating targets for males. After a 30-minute acclimation period, we placed a male on the mating spot with the hermaphrodites and observed male behavior for five minutes. The mating behavior scoring included recording the time stamp of the male’s response to hermaphrodite contact, the number of vulva passes during scanning, and the location of vulva event. We assessed male mating behavior in a blind-to-genotypes manner in three-to-five biological replicates (20 animals per genotype in each replicate). Statistical analysis was conducted using Kruskal-Wallis test.

### Genotyping of alleles used in the study

One-to-four adult hermaphrodites were lysed in 10 μl of the lysis buffer of the following composition: 10 mM Tris pH 8.3, 50 mM KCl, 2.5 mM MgCl_2_, 0.45% NP40, 0.01% w/v gelatin, proteinase K New England Biolabs #P8107 2.5 units/mL (1::300 diluted). Lysis was conducted for 1 hour at 65°C, followed by inactivation of proteinase K at 95°C for 15 min. Final lysates were diluted with water up to 100 μl of total volume, and 4 μl of the diluted lysate was used in a genotyping PCR reaction. Genotyping was performed using Taq polymerase (New England Biolabs #M0273) according to the manufacturer directions with the following PCR cycling parameters: Step1 – 95°C – 2 min, Step2 – 95°C – 20 sec, Step 3 – t_annealing_ – 20 sec, Step 4 – t_elongation_ – 1 min/kb. For specific allele PCR conditions, refer to the table below.

**Table S3:**
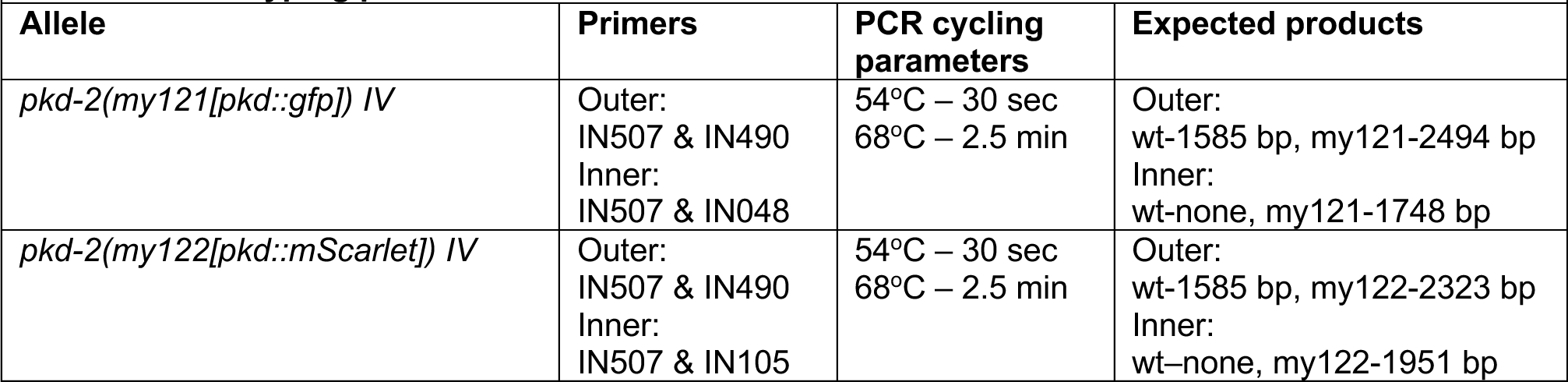
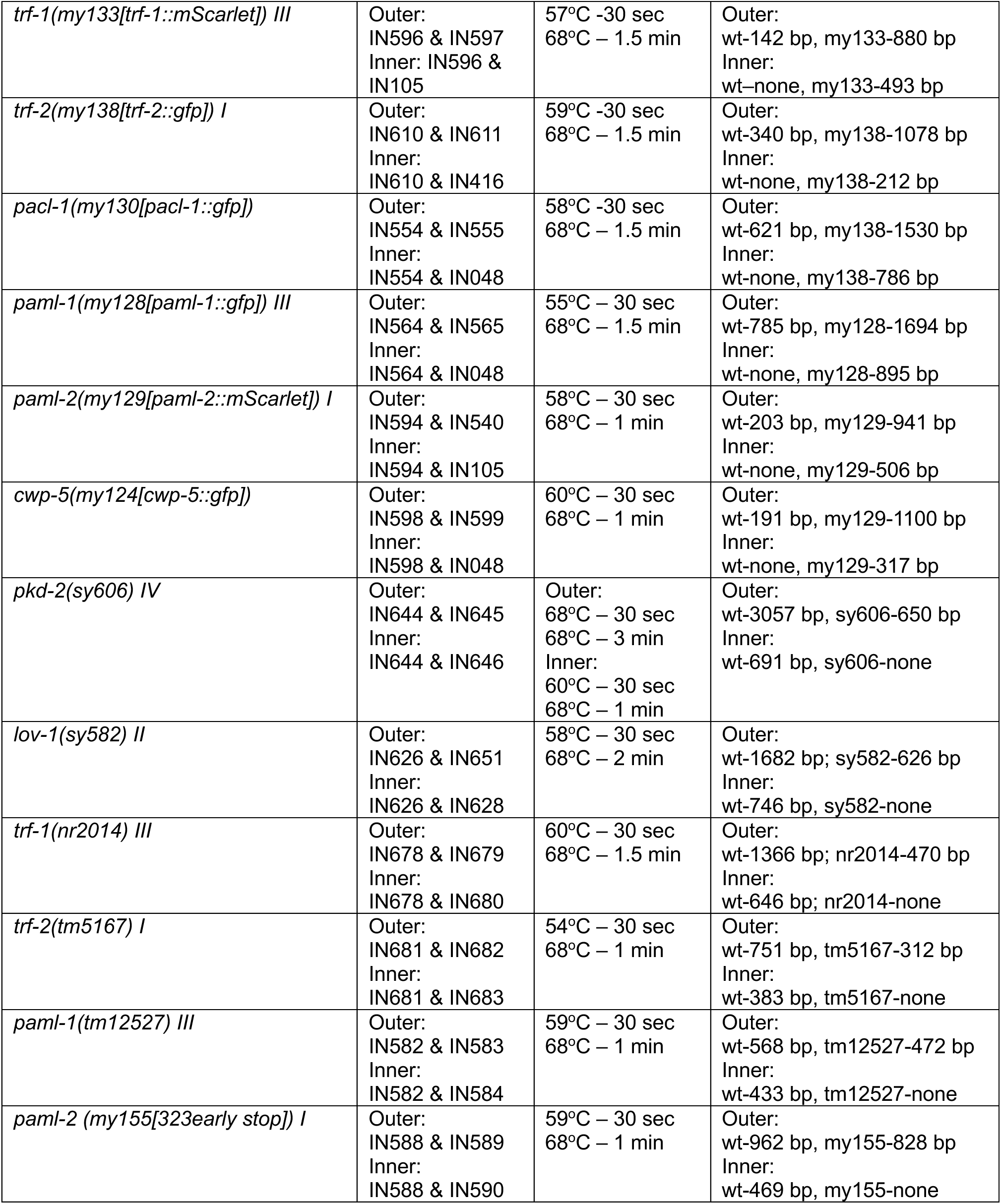
Genotyping parameters.

**Table S4:**
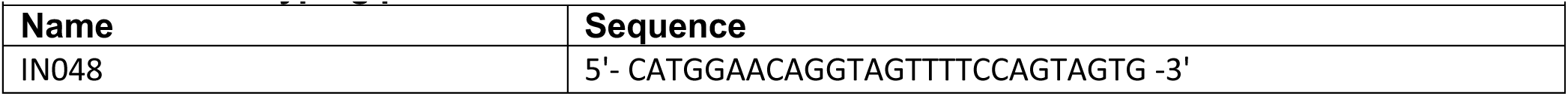
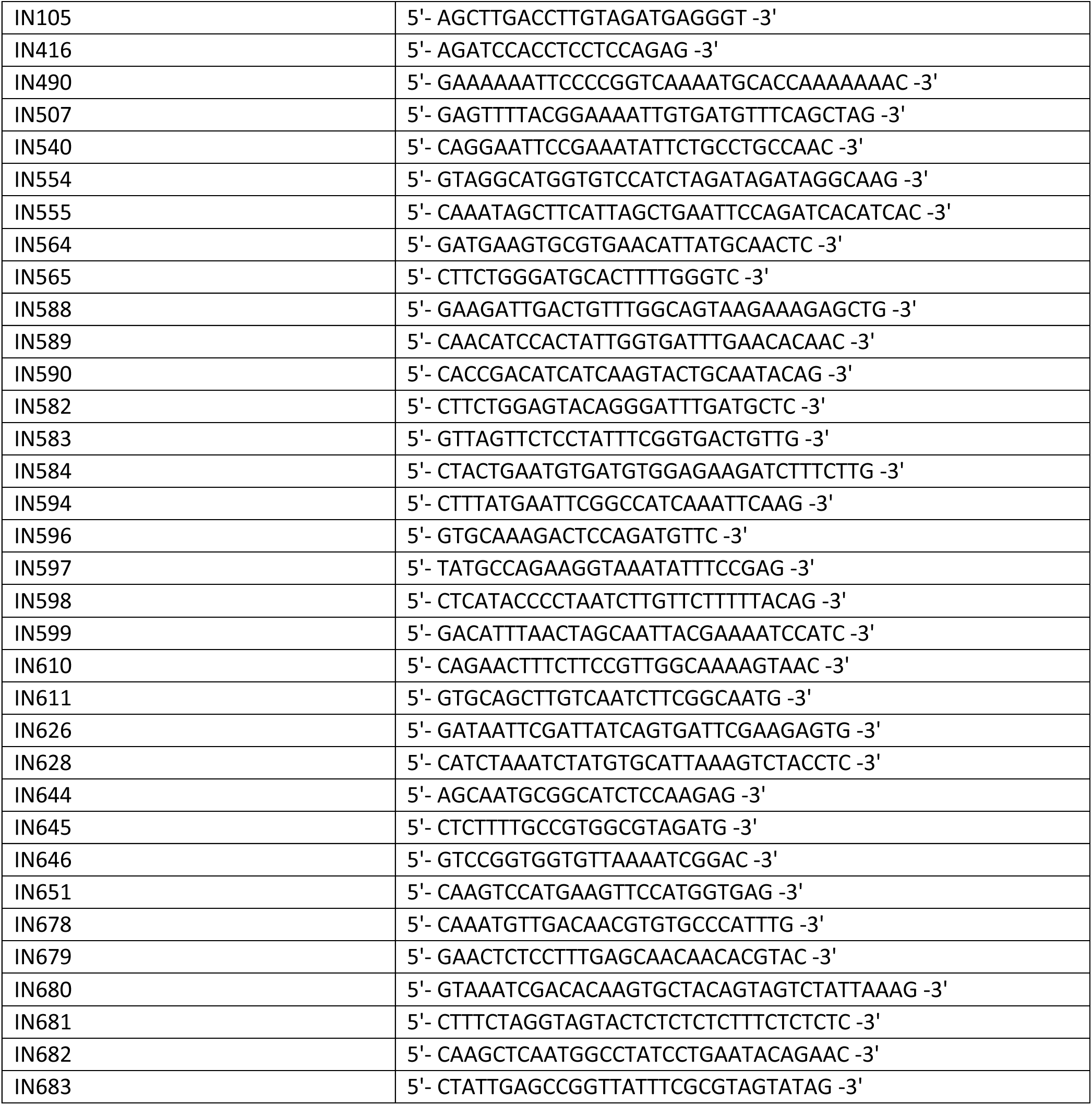
Genotyping primers.

**Table S5:**
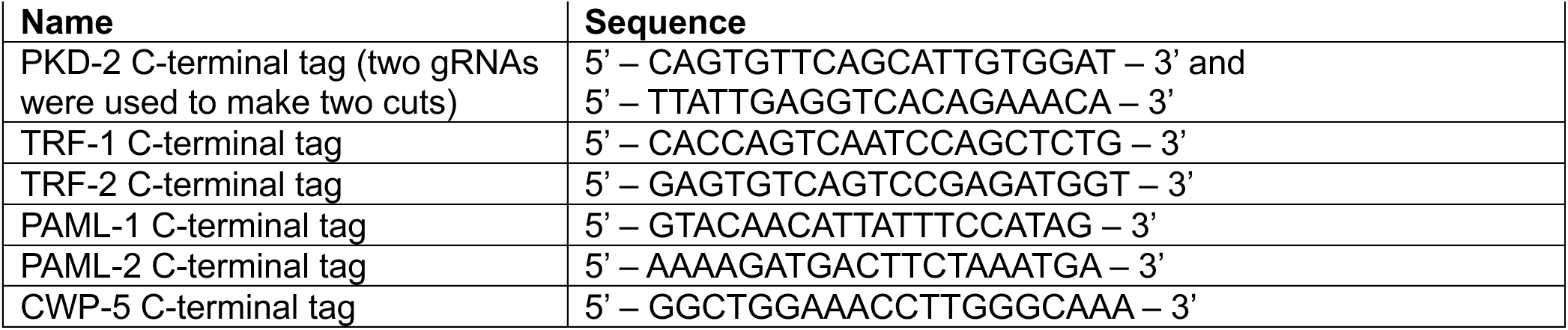
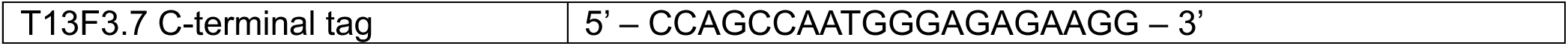
gRNAs for Cas9-mediated genome editing.

### Sequences of endogenous reporters generated in the study

Yellow and orange shading indicate alternating exons.

Grey shading of capital letters indicates encoded flexible linker (GGGGSx4). Green and magenta shading indicate encoded GFP and mScarlet, respectively.

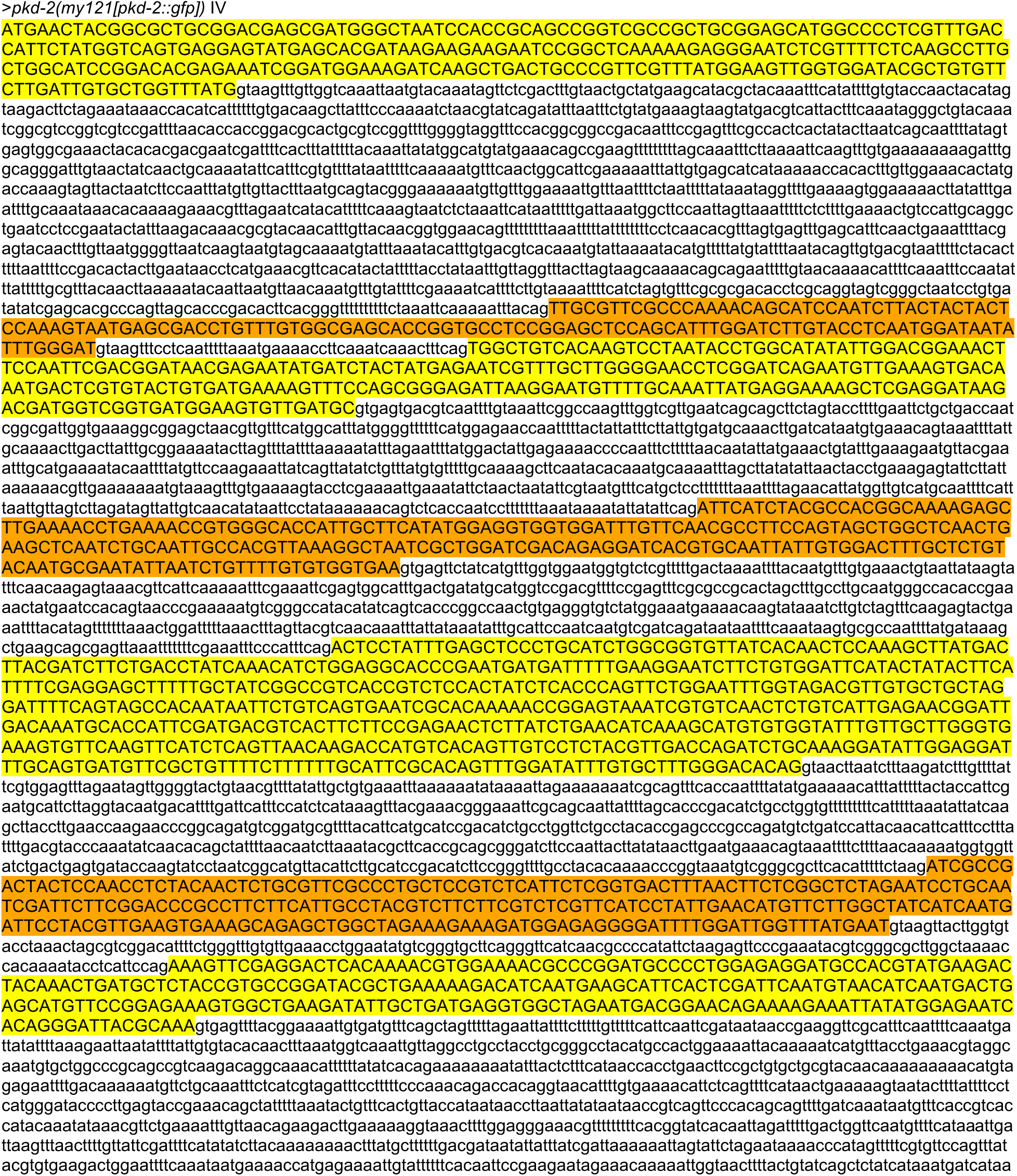
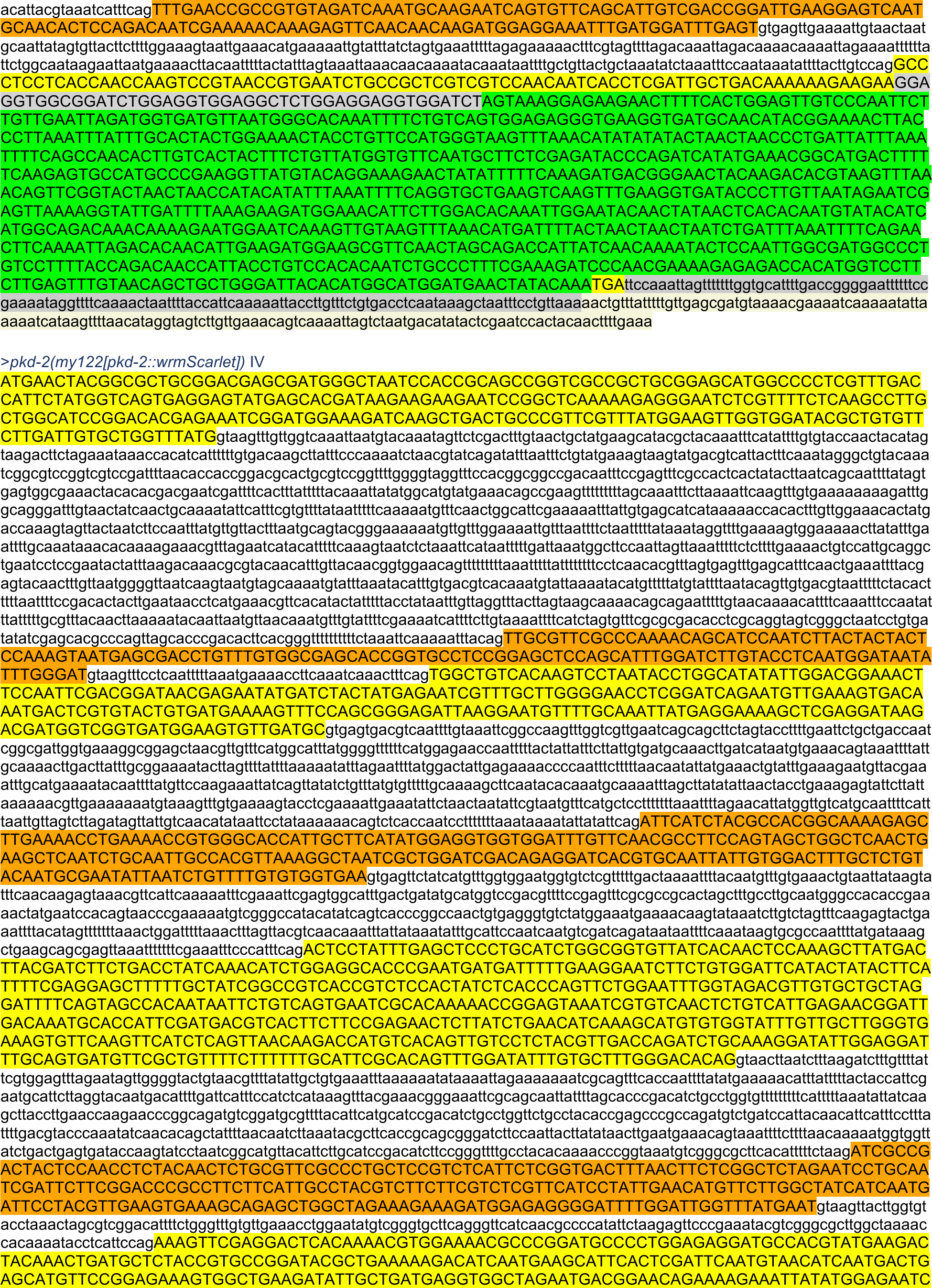
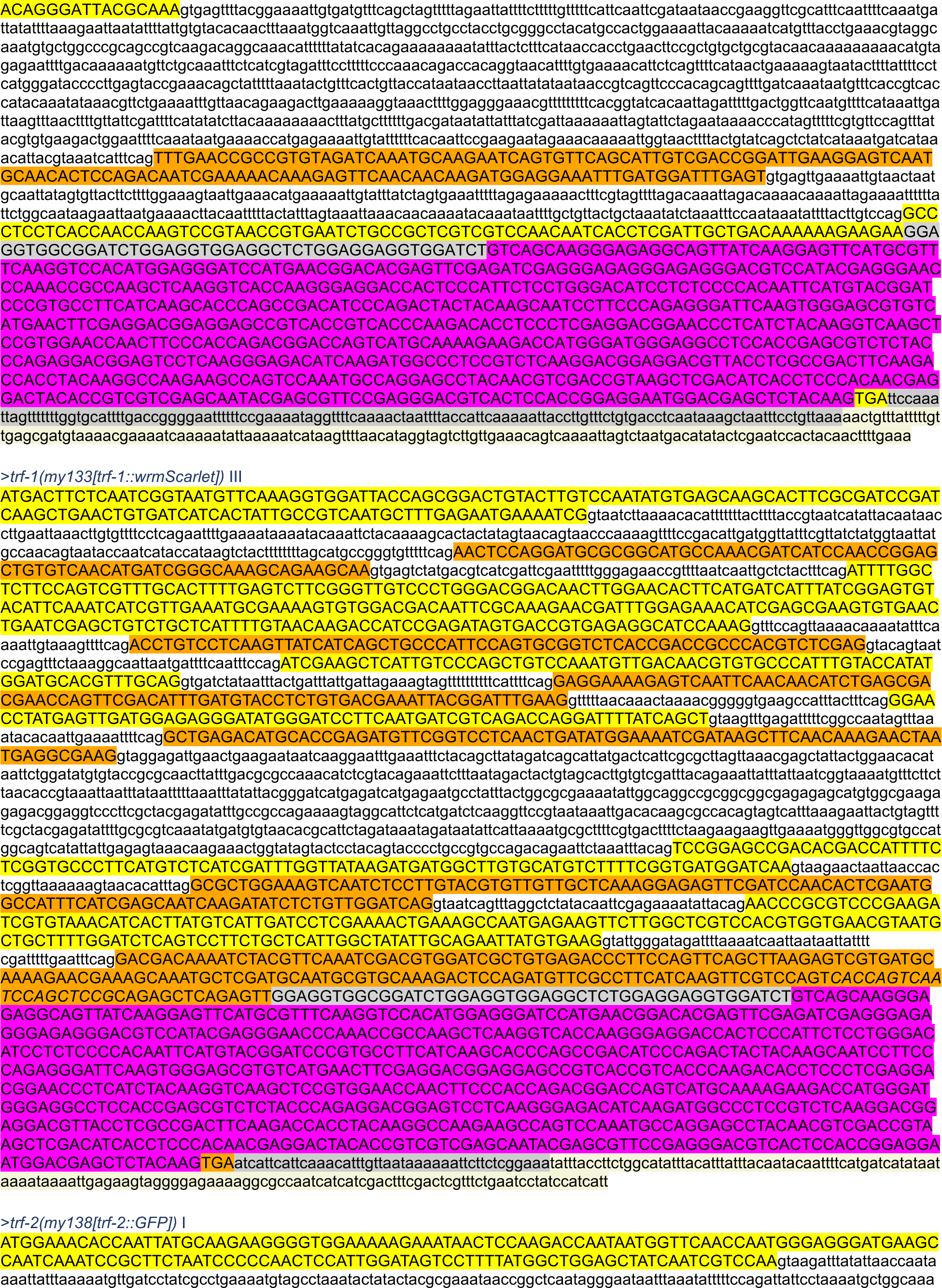
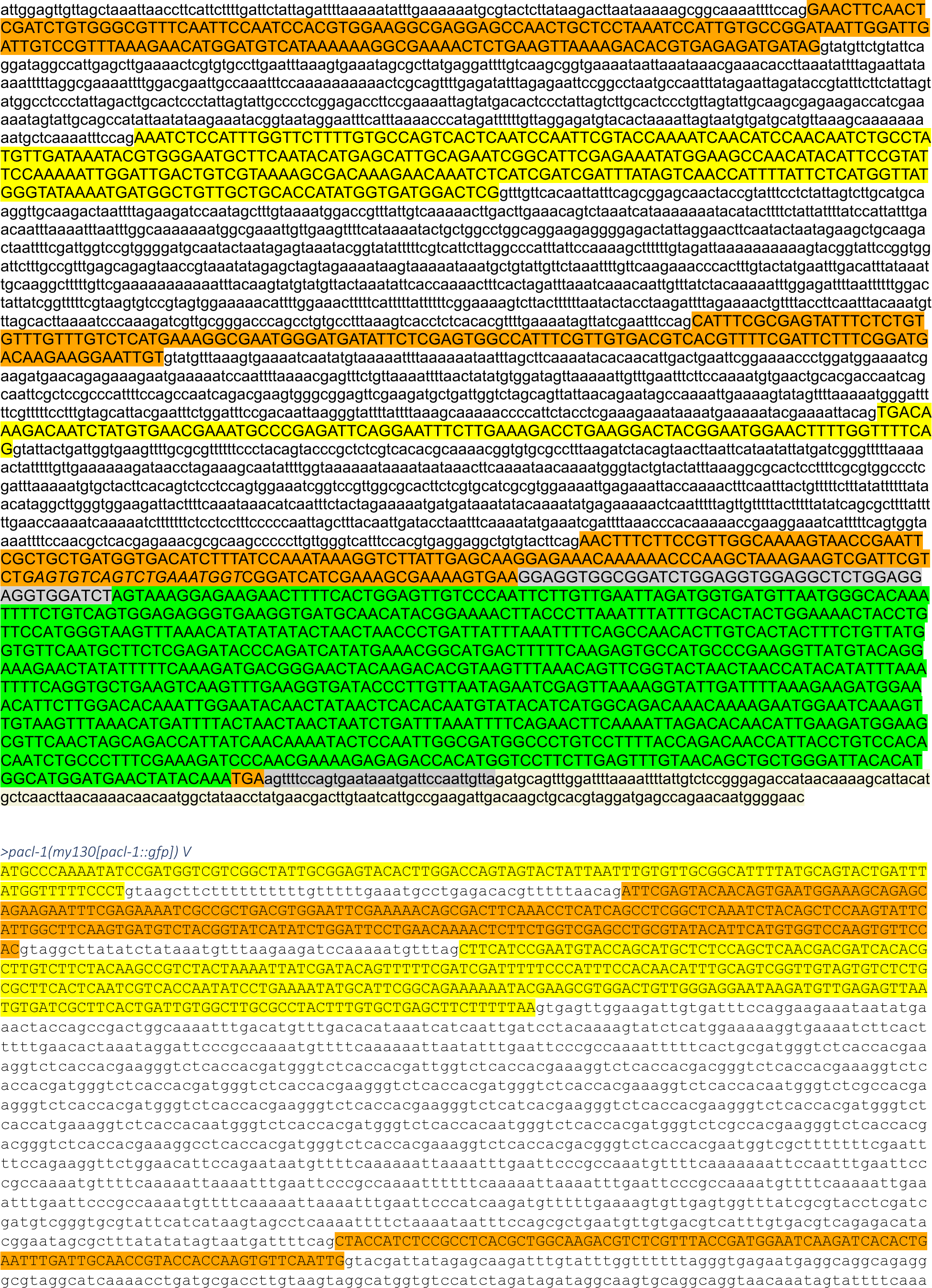
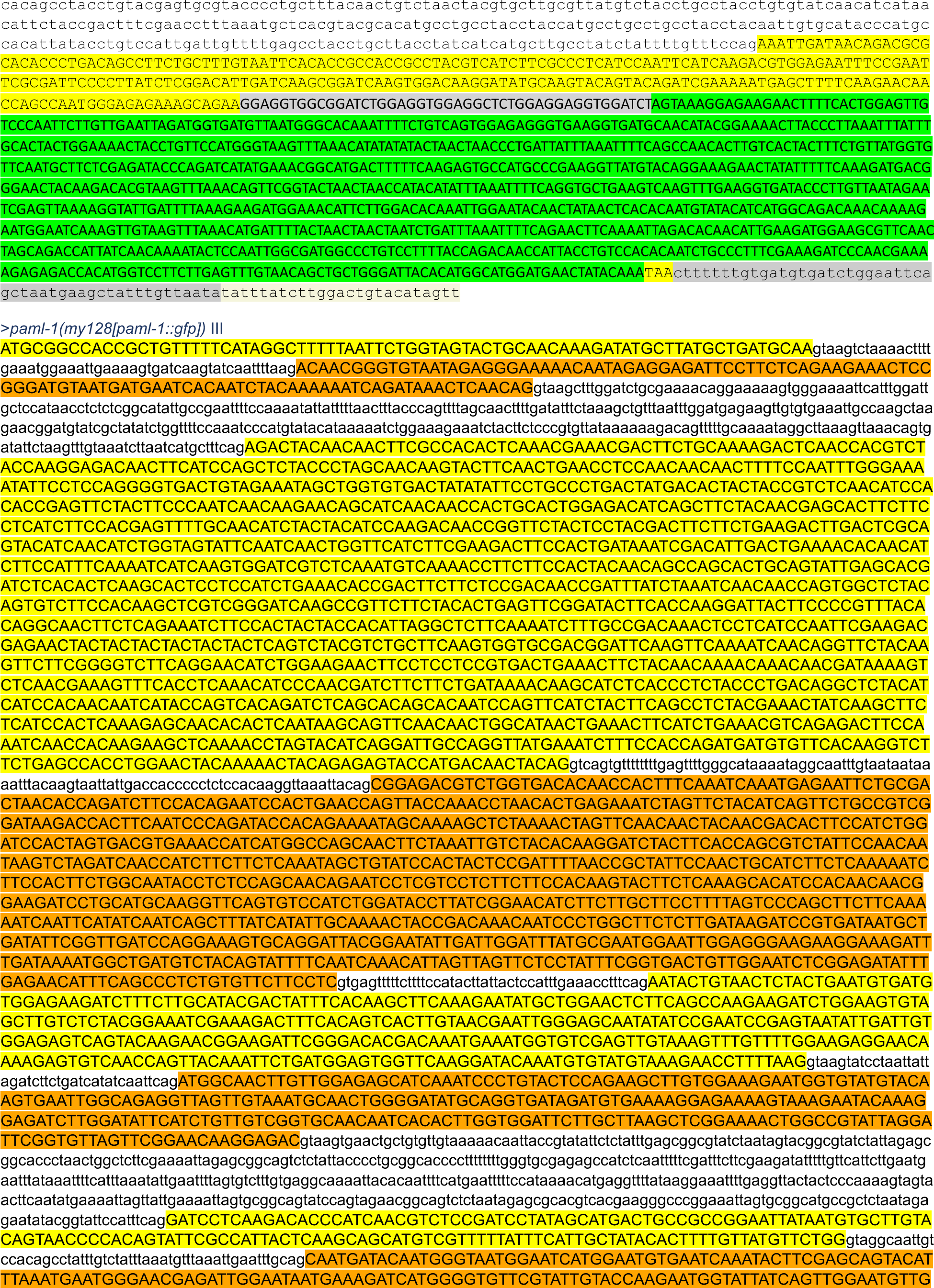
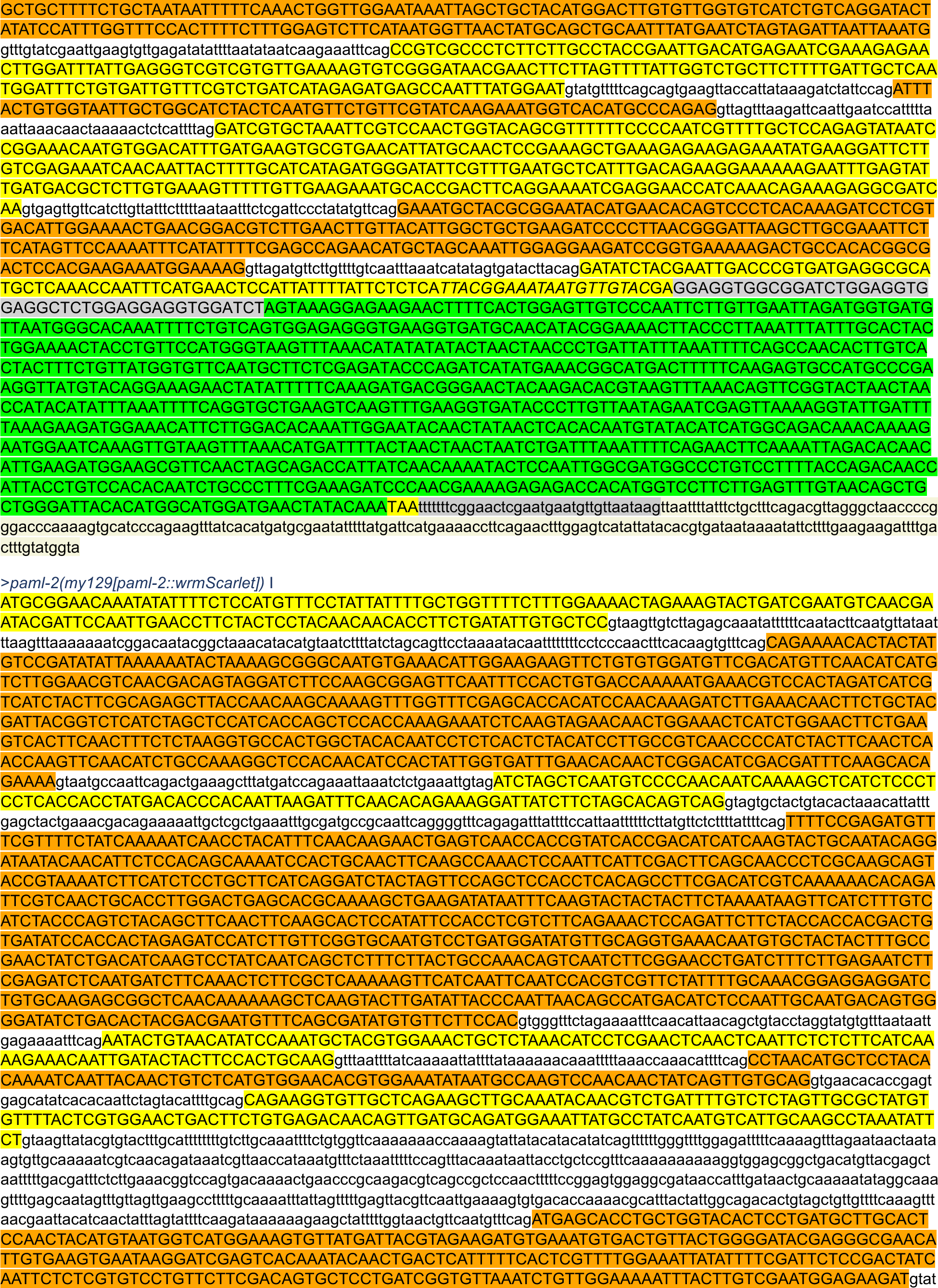
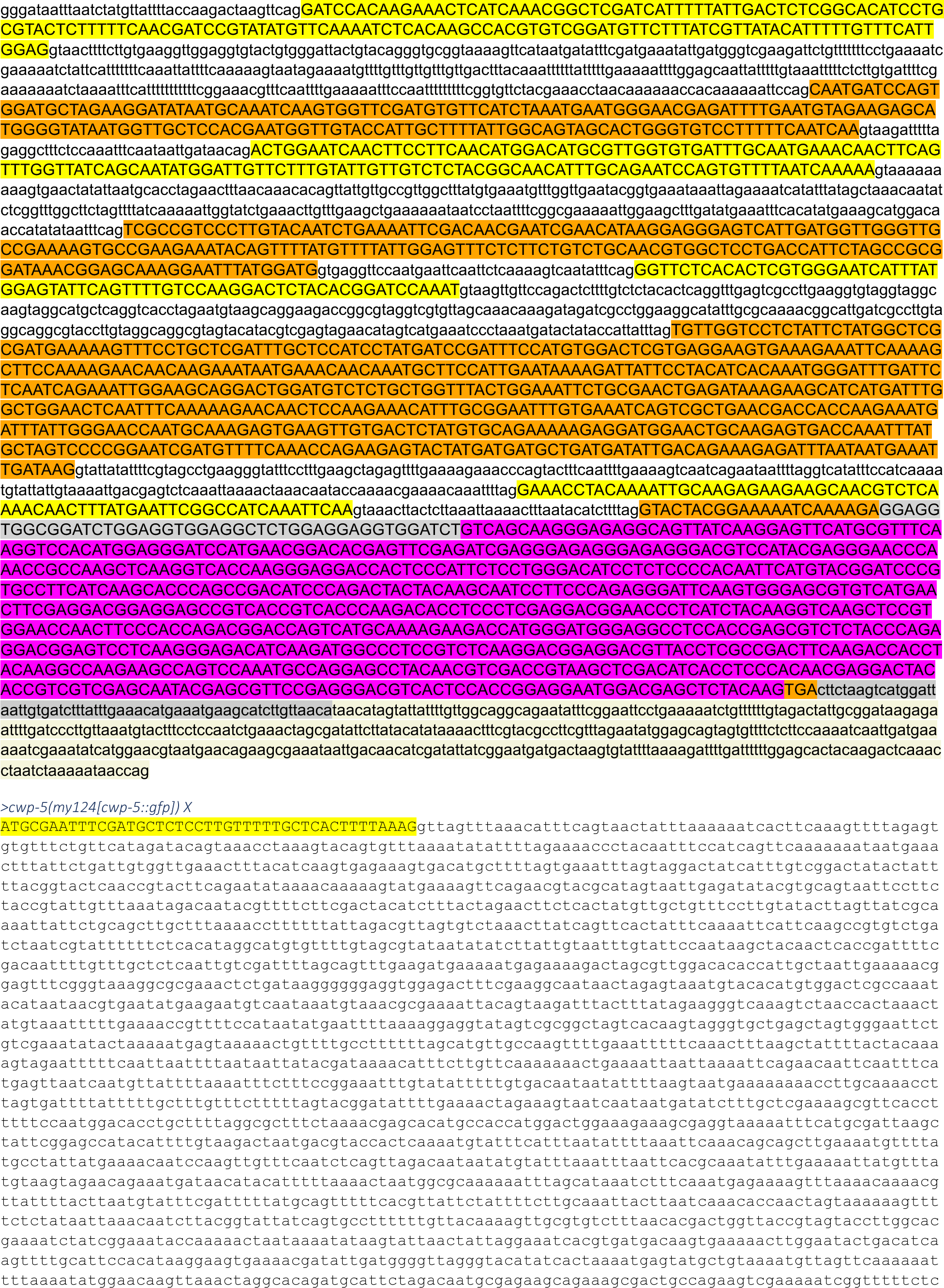
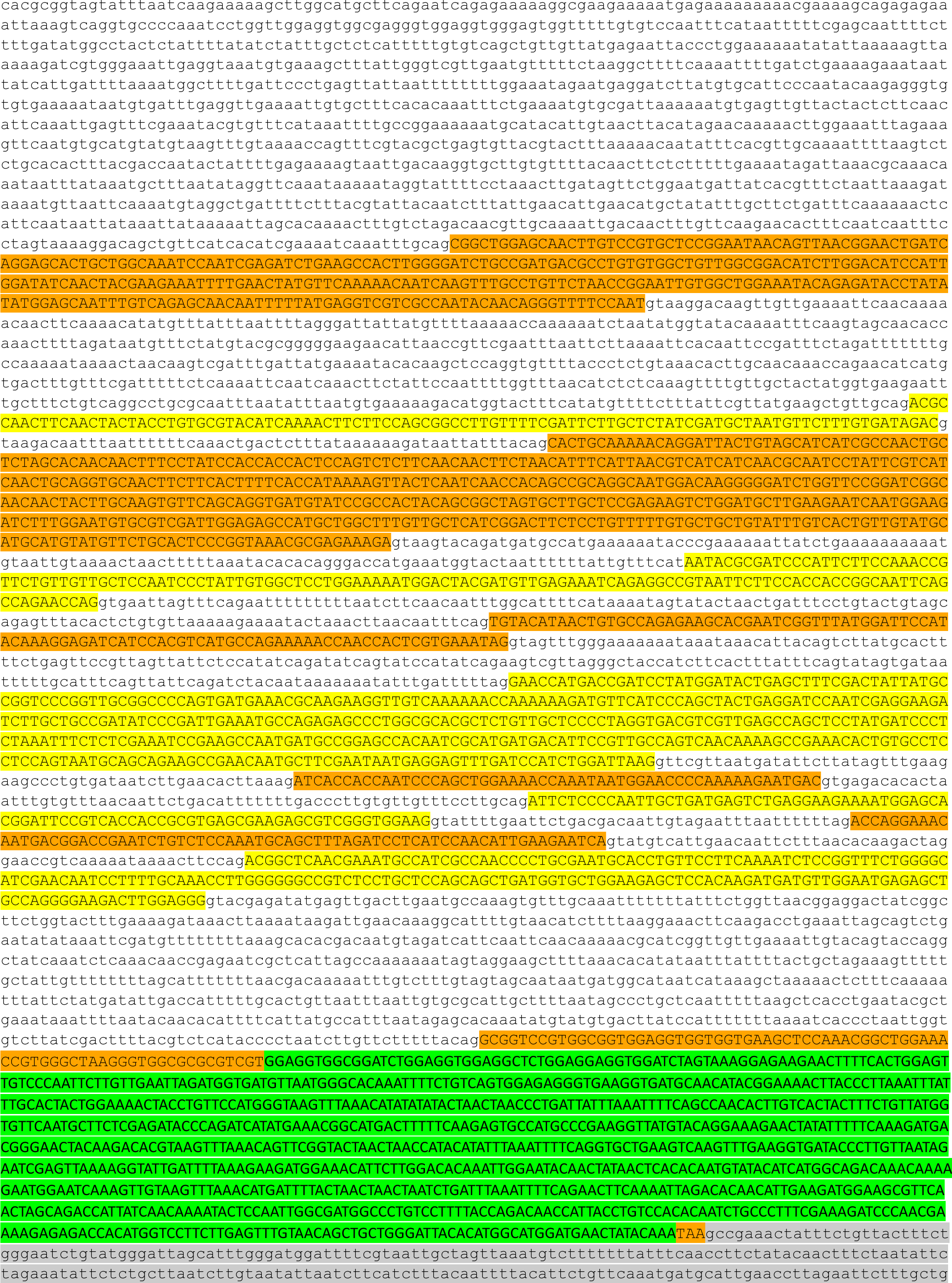
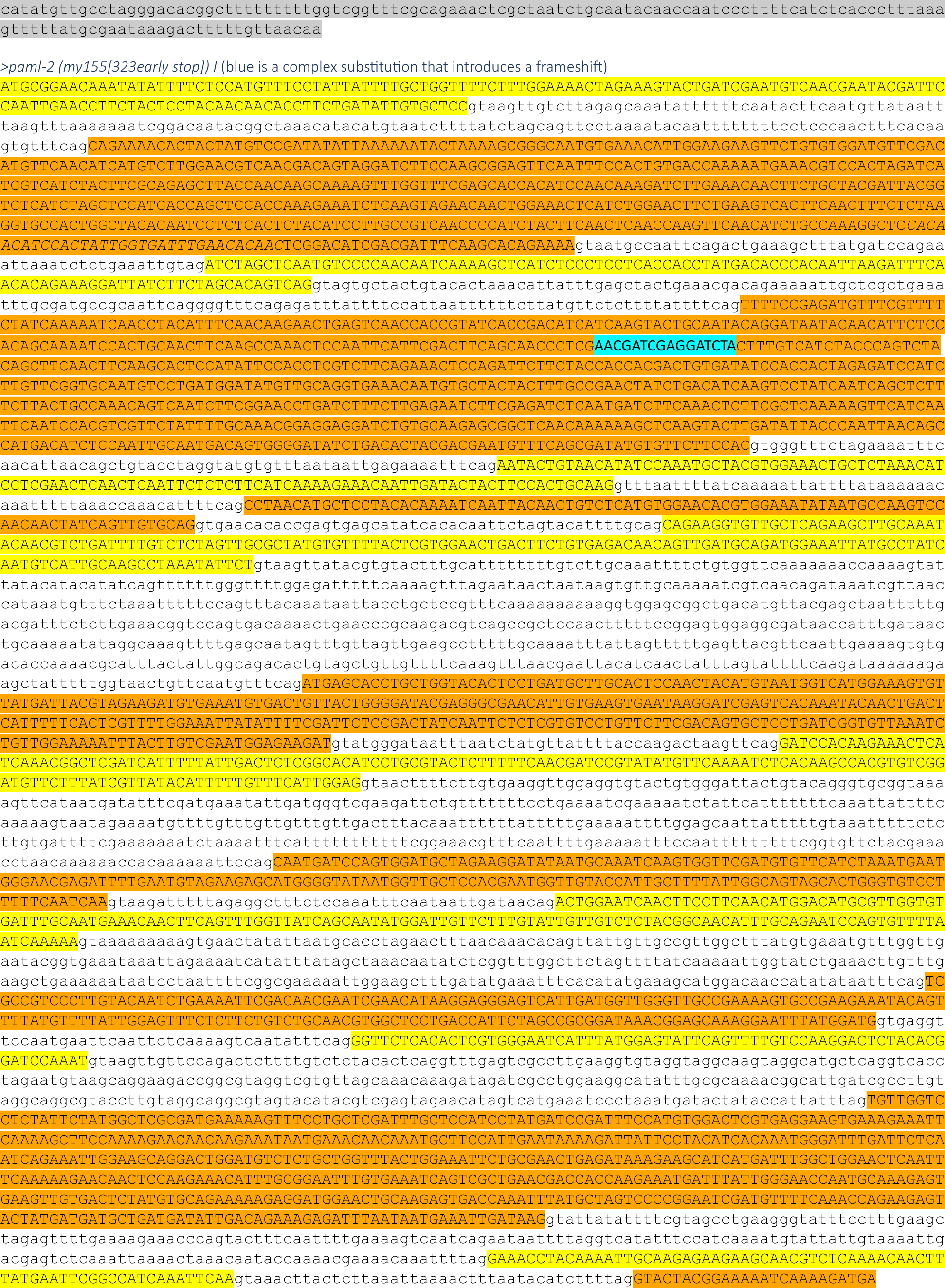
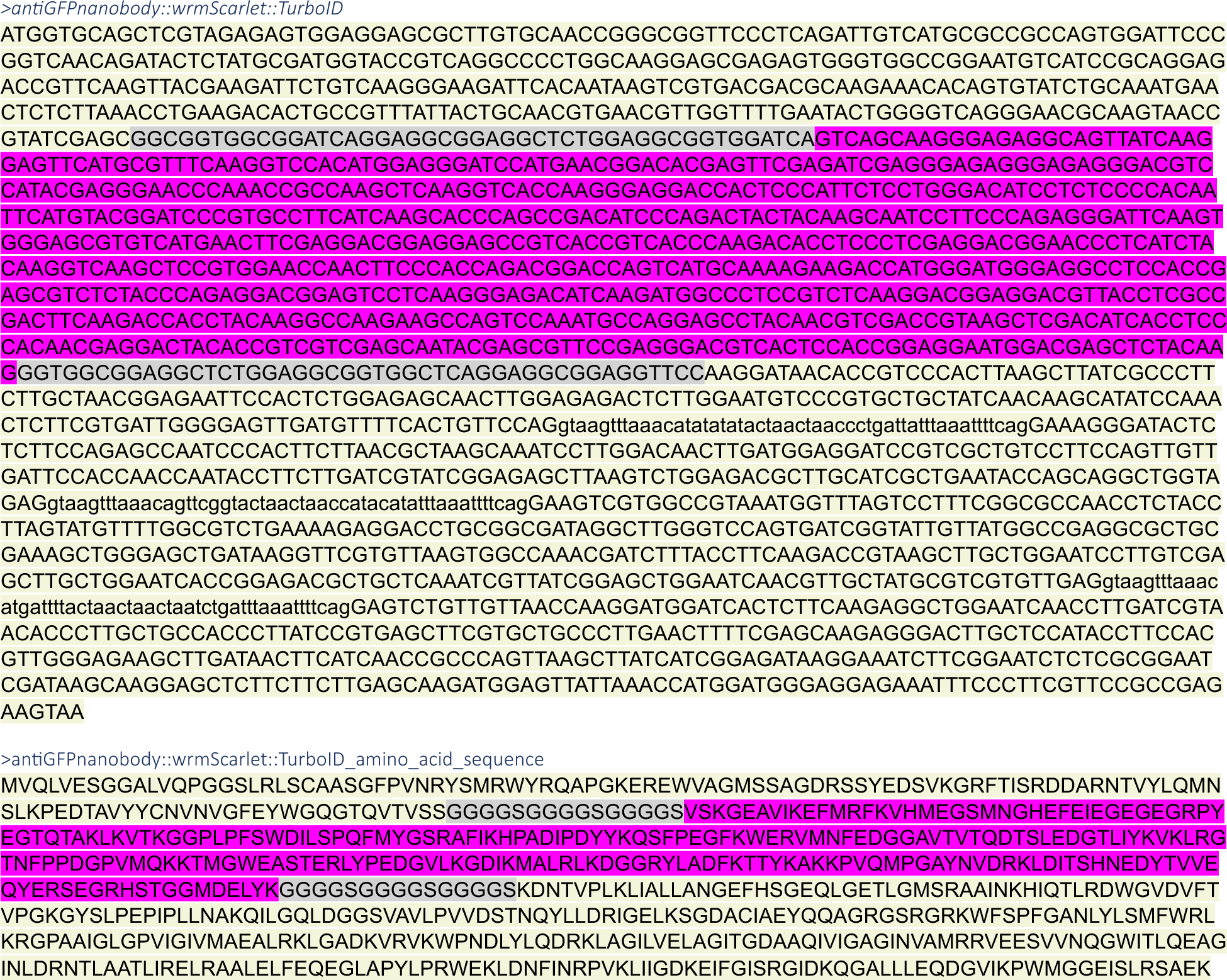

## References

1. Hilgendorf, K. I., Myers, B. R. & Reiter, J. F. Emerging mechanistic understanding of cilia function in cellular signalling. Nat Rev Mol Cell Biol 1–19 (2024).

2. Hogan, M. C. et al. Characterization of PKD protein-positive exosome-like vesicles. J Am Soc Nephrol 20, 278–288 (2009).

3. Wyart, C., Carbo-Tano, M., Cantaut-Belarif, Y., Orts-Del’Immagine, A. & Böhm, U. L. Cerebrospinal fluid-contacting neurons: multimodal cells with diverse roles in the CNS. Nat Rev Neurosci 24, 540–556 (2023).

4. Orts-Del’Immagine, A. et al. Sensory Neurons Contacting the Cerebrospinal Fluid Require the Reissner Fiber to Detect Spinal Curvature In Vivo. Current Biology 30, 827–839.e4 (2020).

5. Djenoune, L. et al. The dual developmental origin of spinal cerebrospinal fluid-contacting neurons gives rise to distinct functional subtypes. Sci Rep 7, 719 (2017).

6. Tanaka, Y., Morozumi, A. & Hirokawa, N. Nodal flow transfers polycystin to determine mouse left-right asymmetry. Developmental Cell (2023) doi:10.1016/j.devcel.2023.06.002.

7. Barr, M. M. & Sternberg, P. W. A polycystic kidney-disease gene homologue required for male mating behaviour in C. elegans. Nature 401, 386–389 (1999).

8. Barr, M. M. et al. The Caenorhabditis elegans autosomal dominant polycystic kidney disease gene homologs lov-1 and pkd-2 act in the same pathway. Curr Biol 11, 1341–1346 (2001).

9. Walsh, J. D. et al. Tracking N– and C-termini of C. elegans polycystin-1 reveals their distinct targeting requirements and functions in cilia and extracellular vesicles. PLoS Genet 18, e1010560 (2022).

10. Barrios, A., Nurrish, S. & Emmons, S. W. Sensory regulation of C. elegans male mate-searching behavior. Curr Biol 18, 1865–1871 (2008).

11. Barrios, A., Ghosh, R., Fang, C., Emmons, S. W. & Barr, M. M. PDF-1 neuropeptide signaling modulates a neural circuit for mate-searching behavior in C. elegans. Nat Neurosci 15, 1675–1682 (2012).

12. Wang, J. et al. C. elegans ciliated sensory neurons release extracellular vesicles that function in animal communication. Curr Biol 24, 519–525 (2014).

13. Wang, J., Nikonorova, I. A., Gu, A., Sternberg, P. W. & Barr, M. M. Release and targeting of polycystin-2-carrying ciliary extracellular vesicles. Curr Biol 30, R755–R756 (2020).

14. Wang, J. et al. Sensory cilia act as a specialized venue for regulated extracellular vesicle biogenesis and signaling. Curr Biol 31, 3943–3951.e3 (2021).

15. Long, H. et al. Comparative Analysis of Ciliary Membranes and Ectosomes. Curr Biol 26, 3327–3335 (2016).

16. Nikonorova, I. A. et al. Isolation, profiling, and tracking of extracellular vesicle cargo in Caenorhabditis elegans. Curr Biol 32, 1924–1936.e6 (2022).

17. Branon, T. C. et al. Efficient proximity labeling in living cells and organisms with TurboID. Nat Biotechnol 36, 880–887 (2018).

18. Holzer, E., Rumpf-Kienzl, C., Falk, S. & Dammermann, A. A modified TurboID approach identifies tissue-specific centriolar components in C. elegans. PLoS Genet 18, e1010150 (2022).

19. Maguire, J. E. et al. Myristoylated CIL-7 regulates ciliary extracellular vesicle biogenesis. Mol Biol Cell 26, 2823–2832 (2015).

20. Wang, J. et al. Cell-Specific Transcriptional Profiling of Ciliated Sensory Neurons Reveals Regulators of Behavior and Extracellular Vesicle Biogenesis. Curr Biol 25, 3232– 3238 (2015).

21. Portman, D. S. & Emmons, S. W. Identification of C. elegans sensory ray genes using whole-genome expression profiling. Dev Biol 270, 499–512 (2004).

22. Miller, R. M. & Portman, D. S. A latent capacity of the C. elegans polycystins to disrupt sensory transduction is repressed by the single-pass ciliary membrane protein CWP-5. Dis Model Mech 3, 441–450 (2010).

23. Park, H. H. Structure of TRAF Family: Current Understanding of Receptor Recognition. Front. Immunol. 9, (2018).

24. Das, A. et al. The Structure and Ubiquitin Binding Properties of TRAF RING Heterodimers. J Mol Biol 433, 166844 (2021).

25. Yin, Q. et al. E2 interaction and dimerization in the crystal structure of TRAF6. Nat Struct Mol Biol 16, 658–666 (2009).

26. Das, A., Foglizzo, M., Padala, P., Zhu, J. & Day, C. L. TRAF trimers form immune signalling networks via RING domain dimerization. FEBS Letters 597, 1213–1224 (2023).

27. Raykov, L., Mottet, M., Nitschke, J. & Soldati, T. A TRAF-like E3 ubiquitin ligase TrafE coordinates ESCRT and autophagy in endolysosomal damage response and cell-autonomous immunity to Mycobacterium marinum. Elife 12, e85727 (2023).

28. Liu, K. S. & Sternberg, P. W. Sensory regulation of male mating behavior in Caenorhabditis elegans. Neuron 14, 79–89 (1995).

29. Ding, H., Li, L. X., Harris, P. C., Yang, J. & Li, X. Extracellular vesicles and exosomes generated from cystic renal epithelial cells promote cyst growth in autosomal dominant polycystic kidney disease. Nat Commun 12, 4548 (2021).

30. Fencl, F. et al. Genotype-phenotype correlation in children with autosomal dominant polycystic kidney disease. Pediatr Nephrol 24, 983–989 (2009).

31. Ha, K. et al. The heteromeric PC-1/PC-2 polycystin complex is activated by the PC-1 N-terminus. Elife 9, e60684 (2020).

32. Lea, W. A. et al. Analysis of the polycystin complex (PCC) in human urinary exosome-like vesicles (ELVs). Sci Rep 10, 1500 (2020).

33. Naito, A. et al. TRAF6-deficient mice display hypohidrotic ectodermal dysplasia. Proceedings of the National Academy of Sciences 99, 8766–8771 (2002).

34. Naito, A. et al. Severe osteopetrosis, defective interleukin-1 signalling and lymph node organogenesis in TRAF6-deficient mice. Genes Cells 4, 353–362 (1999).

35. Lomaga, M. A. et al. TRAF6 deficiency results in osteopetrosis and defective interleukin-1, CD40, and LPS signaling. Genes Dev 13, 1015–1024 (1999).

36. Lomaga, M. A. et al. Tumor necrosis factor receptor-associated factor 6 (TRAF6) deficiency results in exencephaly and is required for apoptosis within the developing CNS. J Neurosci 20, 7384–7393 (2000).

37. Peschel, N. et al. Molecular Pathway-Based Classification of Ectodermal Dysplasias: First Five-Yearly Update. Genes (Basel) 13, 2327 (2022).

38. Kantaputra, P. et al. Dental Anomalies in Ciliopathies: Lessons from Patients with BBS2, BBS7, and EVC2 Mutations. Genes (Basel) 14, 84 (2022).

39. Handa, A., Voss, U., Hammarsjö, A., Grigelioniene, G. & Nishimura, G. Skeletal ciliopathies: a pattern recognition approach. Jpn J Radiol 38, 193–206 (2020).

40. Reimer, A., He, Y. & Has, C. Update on Genetic Conditions Affecting the Skin and the Kidneys. Front Pediatr 6, 43 (2018).

41. David, A. et al. Isolated and syndromic brachydactylies: Diagnostic value of hand X-rays. Diagnostic and Interventional Imaging 96, 443–448 (2015).

42. Katsanis, N. Ciliary proteins and exencephaly. Nat Genet 38, 135–136 (2006).

43. Mishra-Gorur, K. et al. Pleiotropic role of TRAF7 in skull-base meningiomas and congenital heart disease. Proc Natl Acad Sci U S A 120, e2214997120 (2023).

44. Jouret, F. & Devuyst, O. Targeting chloride transport in autosomal dominant polycystic kidney disease. Cellular Signalling 73, 109703 (2020).

45. Link, C. D., Silverman, M. A., Breen, M., Watt, K. E. & Dames, S. A. Characterization of Caenorhabditis elegans lectin-binding mutants. Genetics 131, 867–881 (1992).

46. Davis, P. et al. WormBase in 2022—data, processes, and tools for analyzing Caenorhabditis elegans. Genetics 220, iyac003 (2022).

47. Dokshin, G. A., Ghanta, K. S., Piscopo, K. M. & Mello, C. C. Robust Genome Editing with Short Single-Stranded and Long, Partially Single-Stranded DNA Donors in Caenorhabditis elegans. Genetics 210, 781–787 (2018).

